## Supplementary material for "A flexible empirical Bayes approach to multivariate multiple regression, and its improved accuracy in predicting multi-tissue gene expression from genotypes": S1 Text

### A. Conventions used in mathematical expressions

For the mathematical expressions below, we denote matrices as bold, uppercase letters (e.g., $\boldsymbol{A}$), column vectors as bold, lowercase letters ($\boldsymbol{a}$), and scalars are written in plain, lowercase letters ($a$). We use $\mathbb{R}^{m \times n}$ to denote the set of real $m \times n$ matrices, we use $\boldsymbol{I}_n$ to denote the $n \times n$ identity matrix, $\mathbf{1}_n$ is a column-vector of ones of length $n$, and $\mathrm{tr}(\boldsymbol{A}) = \sum_{i=1}^{n} a_{ii}$ denotes the trace of $n \times n$ matrix $\boldsymbol{A}$.

For describing Bayesian calculations, our convention is to use the "0" subscript to indicate priors and the "1" subscript to indicate posteriors.

### B. Preparation of GTEx data

Here we describe the steps that were taken to prepare the GTEx gene expression and genotype data for *mr.mash*. These same steps were taken for both the application and the simulations, but note that only the genotype data were used in the simulations.

The GTEx v8 data were used for all the analyses. Files containing expression data (RNASeq QC version 1.1.9 TPM), covariate data and genotypes (VCF format) were downloaded from dbGaP, study accession phs000424.v8.p2. The raw expression and genotype data were preprocessed using data processing scripts available online. The data processing scripts used are available at: `https://github.com/gaow/mvarbvs/blob/master/analysis/gtex-v7/20170929_Expression_Covariate.ipynb`, `https://github.com/gaow/mvarbvs/blob/master/analysis/gtex-v7/20170929_Genotype.ipynb`, and `https://github.com/gaow/mvarbvs/blob/master/workflow/GTEx_V8_preprocessing.ipynb`. All these scripts are intended to be run using the SoS (Script of Scripts) framework available at `https://github.com/vatlab/sos`. These scripts were all run with the default parameters, e.g., `sos run GTEx_V8_preprocessing.ipynb`. These scripts were used to extract and reformat the genotype and expression data from the GTEx v8 data files for analysis with *mr.mash* and the other methods. Data processing steps included reformatting the data, removing the effects of selected covariates from the gene expression levels, and selecting biallelic genetic variants 1 Mb upstream and downstream of the transcription start site (TSS) for each gene. The result of running this pipeline was an R object (`.rds`) file for all genes (including pseudogenes) on chromosomes 1–22. In total, 39,840 `.rds` files were generated. Each file contained an $n \times p$ genotype matrix $\boldsymbol{X}$ and an $n \times r$ matrix of gene expression residuals $\boldsymbol{Y}$, possibly containing missing values. For a given gene, the number of samples ($n$) was at most 848, and the number of tissues ($r$) was at most 49. (Data preparation done as part of the GTEx project included filtering out tissues for each gene (see `https://gtexportal.org/home/methods` for details), so $r$ was often less than 49 for a given gene.) We also removed samples in which expression measurements were missing in all tissues.

After running this data processing pipeline, the following additional filtering steps were applied, separately for each gene, to the genotype data $\boldsymbol{X}$ before running *mr.mash* or one of the other multivariate regression methods. Genetic variants satisfying these conditions were retained: genetic variants with minor allele frequency (MAF) greater than 0.05 (these are MAFs calculated from the GTEx genotypes); missing genotype rate less than 0.05; and genotype variance greater than 0.05. For any genetic variants that had one or more missing genotypes, the missing genotypes were filled in using the mean genotype for that variant.

After taking these steps, on average there were 4,465 genetic variants for a given gene, but this number varied widely from gene to gene, from 41 to 21,247. The number of tissues, $r$, varied from 1 to 49, with a median of 43. The number of samples, $n$, varied from 73 to 838, with a median of 836. The rate of missing entries in $\boldsymbol{Y}$ ranged from 0% to 77%, with a median of 63%.

Finally, we removed the "kidney cortex" tissue from $\boldsymbol{Y}$ because it had a maximum sample size less than 100.

### C. Simulations with GTEx genotypes

In brief, for each simulation, we generated multi-tissue expression data sets for 2,000 genes randomly selected from chromosomes 1–22. We only considered genes in which all 838 genotype samples were retained by the data processing pipeline above.

For each gene, we simulated the $n \times r$ gene expression matrix $\boldsymbol{Y}$ for $n = 838$ samples and $r = 10$ tissues using the multivariate multiple regression model. The coefficients $\boldsymbol{B}$ and the residual covariance matrix $\boldsymbol{V}$ used to simulate the phenotypes were chosen so as to capture a variety of genetic architectures: different numbers of genetic variants affecting the phenotypes; different patterns of effect sharing across tissues; and different levels of genomic heritability (including no heritability—that is, simulations in which one or more expression levels were unaffected by genotype). Once we simulated the expression levels, we split the samples into two subsets, one containing 80% of the samples (a training set with $n = 670$) and another containing 20% of the samples (a test set with $n = 168$).

The training data for the 2,000 genes were used to estimate data-driven covariance matrices from the *mr.mash* prior (see Sec. E.4.3). Then we selected 1 out of the 2,000 genes, and fit regression models (*Elastic Net*, *mr.mash*, *etc*) to the genotype and phenotype data from the training set for that gene. Finally, we evaluated the accuracy of each model's predictions in the test samples for that same gene.

#### *C.1. Simulation scenarios*

The five simulation scenarios are described briefly in the main text, and here we describe them in greater detail.

A. In the "Equal Effects" scenario, a causal variant affected all tissues, and the effects were exactly the same in all tissues. This was the setting in which we had most to gain by analyzing all tissues simultaneously compared to analyzing the tissues one at a time.

B. In the "Independent Effects" scenario, a causal variant affected all tissues, and the effects were independent across all tissues (more precisely, the effects were independent conditioned on the genetic variant being a causal variant). The Independent Effects scenario represented the situation in which there was less to gain from a joint analysis of all tissues because the effect sizes were not shared across tissues. Nonetheless, we expected some gain in a multivariate analysis compared to a tissue-by-tissue analysis because knowing that a genetic variant has an effect in one tissue was helpful for detecting an effect in other tissues.

C. In the "Mostly Null" scenario, a causal variant only affected the first tissue, and therefore the remaining tissues were unaffected by genotype. This represented a scenario in which the effects for the given gene were tissue-specific. We did not expect any gain in performing a joint analysis of all tissues in this setting because there was no sharing of effects across tissues. In fact, there was the potential of multivariate analysis methods to harm performance by incorrectly shrinking effect estimates for the first tissue toward effect estimates in the remaining "null" tissues.

D. In the "Equal Effects + Null" scenario, the effects were equal across tissue 1 through 3, and there were no effects in tissues 4 through 10.

E. In the "Shared Effects in Subgroups" scenario, effects were drawn from a mixture of effect-sharing patterns: 50% of the time, the effects were shared (unequally) across tissues 1 through 3 and explained 20% of the variance of each tissue; in the other 50% of the time, the effects are shared (unequally) in tissues 4 through 10 and explained only 5% of the variance of each tissue. This scenario was intended to correspond roughly to the patterns of effect sharing estimated from previous multivariate analyses of GTEx Project data (see for example Fig. 3a in Urbut et al. 2019).

In the first four scenarios, the genetic effects explain 20% of the variance of each tissue. (Except in the Mostly Null scenario, where the genetic effects explained none of the variance in all tissues other than the first, and in the Equal Effects + Null scenario, where the genetic effects explained none of the variance in tissues 4 through 10.)

#### C.2. Simulation procedure

For all simulation scenarios, we used the following procedure to simulate the multi-tissue expression. For all simulations, $n = 838$ and $r = 10$.

1. Choose a gene at random.
2. Center the columns of the $n \times p$ genotype matrix $\boldsymbol{X}$ so that each column has a mean of zero.
3. Choose the number of causal variants, $p_{\text{causal}}$, uniformly at random among integers 1–10.
4. Choose the $p_{\text{causal}}$ causal variants uniformly at random from $\{1, \ldots, p\}$.
5. Set $\boldsymbol{b}_j = (0, \ldots, 0)^T \in \mathbb{R}^r$ for each non-causal variant $j$.
6. For each causal variant $j$, simulate $\boldsymbol{b}_j \in \mathbb{R}^r$: for the Independent Effects scenario, $\boldsymbol{b}_j \sim N_r(\boldsymbol{0}, \boldsymbol{I}_r)$; for the Equal Effects scenario, $\boldsymbol{b}_j \sim N_r(\boldsymbol{0}, \boldsymbol{1}_r\boldsymbol{1}_r^T)$, where $\boldsymbol{1}_r = (1, \ldots, 1)^T \in \mathbb{R}^r$; for the Mostly Null scenario, $\boldsymbol{b}_j \sim N_r(\boldsymbol{0}, \boldsymbol{S})$, where $\boldsymbol{S}$ is an $r \times r$ matrix of all zeros except for a single one, $s_{11} = 1$; for the Equal Effects + Null scenario, $\boldsymbol{b}_j \sim N_r(\boldsymbol{0}, \boldsymbol{S})$, where $\boldsymbol{S}$ is an $r \times r$ matrix of all zeros except for all ones in the $3 \times 3$ top-right subblock, $s_{ij} = 1$, $i \leq 3, j \leq 3$; for the Shared Effects in Subgroups scenario, the effects $\boldsymbol{b}_j$ were simulated using a mixture of zero-centered multivariate normals. The mixture weights and covariance matrices for this scenario are given in the Zenodo repository Morgante et al. (2023).
7. Form the residual covariance matrix across tissues, $\boldsymbol{V}$. This is a diagonal $r \times r$ matrix in which the diagonal entries $v_{ii}$ are chosen to achieve the target PVE (proportion of variance explained) for each tissue. For tissues $i$ in which genetic variants did not explain variation in expression, we set $v_{ii} = 1$.
8. Simulate the $n \times r$ matrix of gene expression levels, $\boldsymbol{Y} \sim MN(\boldsymbol{B}_0 + \boldsymbol{XB}, \boldsymbol{I}_n, \boldsymbol{V})$, where $\boldsymbol{B}_0$ is an $n \times r$ matrix of ones, $\boldsymbol{B}$ is the $p \times r$ coefficients matrix with rows $\boldsymbol{b}_j$, and $\boldsymbol{I}_n$ is the $n \times n$ identity matrix.
9. Randomly split the data—rows of $\boldsymbol{X}$ and rows of $\boldsymbol{Y}$—into a training set (80% of the data, or 670 individuals) and a test set (20% of the data, or 168 individuals). This training-test split was different for each scenario, but the same for all simulations in a scenario.

**Algorithm 1** Sketch of the mr.mash variational empirical Bayes algorithm allowing for missing data. The matrix $\boldsymbol{Y}$ may contain missing values; $\boldsymbol{Y}_{\text{obs}}$ denotes the set of observed values. The matrix $\boldsymbol{X}$ may be "standardized"—columns scaled to unit standard deviation—which typically reduces computation because more calculations can be reused. In the algorithm, $\tilde{\boldsymbol{x}}_j$ denotes the $j$th column of $\tilde{\boldsymbol{X}}$, and $\boldsymbol{b}_j$ is the $j$th row of $\boldsymbol{B}$ (stored as a column vector). "BMSR-mix" is explained in the text. Note this is only an outline of the algorithm, and actual implementation differs to avoid redundant calculations and reduce memory usage.

**Require:** Data inputs $\boldsymbol{X}$ ($n \times p$ matrix) and $\boldsymbol{Y}$ ($n \times r$ matrix).
**Require:** Prior covariances $\mathscr{S}_0 = \{\boldsymbol{S}_{0,1}, \ldots, \boldsymbol{S}_{0,K}\}$.
**Require:** Initial estimates $\boldsymbol{b}_0, \boldsymbol{B}, \boldsymbol{w}_0, \boldsymbol{V}$.
  **repeat**
    Center the columns of $\boldsymbol{X}$ and store the result in $\tilde{\boldsymbol{X}}$.
    If $\boldsymbol{Y}$ contains missing values, impute the missing values using $\boldsymbol{Y}_{\text{obs}}, \boldsymbol{b}_0, \boldsymbol{B}, \boldsymbol{V}$.
    Store the observed and imputed values in $\boldsymbol{Y}_{\text{full}}$.
    Center the columns of $\boldsymbol{Y}_{\text{full}}$ and the store result in $\tilde{\boldsymbol{Y}}$.
    Compute the expected residuals, $\boldsymbol{R} \leftarrow \tilde{\boldsymbol{Y}} - \tilde{\boldsymbol{X}}\boldsymbol{B}$.
    **for** $j$ in $1, \ldots, p$ **do**
      Remove variable $j$ from the expected residuals, $\boldsymbol{R}_j \leftarrow \boldsymbol{R} + \tilde{\boldsymbol{x}}_j \boldsymbol{b}_j^\intercal$.
      $(w_{1,1}^{(j)}, \ldots, w_{1,K}^{(j)}, \boldsymbol{b}_{1,1}^{(j)}, \ldots, \boldsymbol{b}_{1,K}^{(j)}, \boldsymbol{S}_{1,1}^{(j)}, \ldots, \boldsymbol{S}_{1,K}^{(j)}) \leftarrow \text{BMSR-mix}(\boldsymbol{x}_j, \boldsymbol{R}_j, \boldsymbol{V}, \mathscr{S}_0, \boldsymbol{w}_0)$.
      Compute the posterior mean estimates, $\boldsymbol{b}_j = \sum_{k=1}^K w_{1,k}^{(j)} \boldsymbol{b}_{1,k}^{(j)}$.
      Store $\boldsymbol{b}_j$ in $j$th column of $\boldsymbol{B}$.
      Include variable $j$ in the expected residuals, $\boldsymbol{R} \leftarrow \boldsymbol{R}_j - \tilde{\boldsymbol{x}}_j \boldsymbol{b}_j^\intercal$.
    Update the prior, $w_{0,k} \leftarrow \sum_{j=1}^p w_{1,k}^{(j)}/p$.
    Update the residual covariance matrix, $\boldsymbol{V} \leftarrow E_q[(\boldsymbol{Y}_{\text{full}} - \tilde{\boldsymbol{X}}\boldsymbol{B})^T(\boldsymbol{Y}_{\text{full}} - \tilde{\boldsymbol{X}}\boldsymbol{B})]/n$.
    Update the intercept, $\boldsymbol{b}_0 \leftarrow (\boldsymbol{Y}_{\text{full}} - \boldsymbol{X}\boldsymbol{B})^\intercal \boldsymbol{1}_n/n$.
  **until** convergence criterion is met or maximum iteration is reached
  **return** Intercept $\boldsymbol{b}_0$, coefficients $\boldsymbol{B}$, imputed responses $\boldsymbol{Y}_{\text{full}}$, prior weights $\boldsymbol{w}_0$, and residual covariance $\boldsymbol{V}$.

#### C.3. Computing environment

All simulations were run on Linux machines (CentOS Linux 8.4-2105) with Intel Xeon Gold 6248R ("Cascade Lake") processors. We used R 4.0.3 (R Core Team, 2020) linked to OpenBLAS 0.3.12 (conda package r-base 4.0.3, build h349a78a_8 installed via conda-forge). At most 4 GB of memory was needed to run *mr.mash* or one of the other methods on a simulated data set. All methods and other computations were run on a single CPU.

### D. Algorithms for fitting the *mr.mash* model

An outline of the variational empirical Bayes algorithm for *mr.mash* is given in Algorithm 1. This algorithm also handles imputation of missing data which we describe in later sections. Algorithm 2 gives a more detailed description of the *mr.mash* algorithm with no missing data. The algorithm for $\boldsymbol{Y}$ with missing values is given in Algorithm 3.

A brief derivation of Algorithm 2 is given in the next two subsections, and more detailed derivations are given in Sections F and G. The algorithm for *mr.mash* with missing data is derived in Sec. H.

The steps of Algorithm 2 that involve the most effort are the updates to the posteriors $q_j(\boldsymbol{b}_j)$ in the inner loop over $j$, and the update to $\boldsymbol{V}$. The update to $\boldsymbol{V}$ involves computing the ERSS (13), which involves $O((n+p)r^2)$ flops. Each update to $q_j(\boldsymbol{b}_j)$ involves computing $\bar{\boldsymbol{R}}_j$ and the BMSR-mix posterior, requiring $O(nr + kr^3)$ flops. Therefore, the computational complexity of a single update of all the parameters (a single iteration of the outer loop) is $O(p(nr + kr^3) + nr^2)$. The effort in the inner-loop updates is greatly reduced when $\boldsymbol{X}$ is standardized because the most expensive computations can be reused (Sec. I). In this special case, the complexity of a single outer-loop iteration reduces to $O(p(nr + kr^2) + nr^2 + kr^3)$. The complexity of the missing data imputation step in Algorithm 3 in

**Algorithm 2** *mr.mash* with no missing data

**Require:** $n \times p$ data matrix $\boldsymbol{X}$ and $n \times r$ data matrix $\boldsymbol{Y}$. Columns of $\boldsymbol{X}$ and $\boldsymbol{Y}$ should be "centered" so that each column has a mean of zero. Optionally, $\boldsymbol{X}$ may also be "standardized"—that is, scaled so that each column has unit standard deviation—which simplifies downstream posterior computations (Sec. I).

**Require:** Set of $K$ covariance matrices, $\mathscr{S}_0$.

**Require:** Initial estimates of the posterior mean coefficients, stored as a $p \times r$ matrix, $\bar{\boldsymbol{B}}$.

**Require:** Initial estimates of the prior mixture weights, $\boldsymbol{w}_0 = (w_{0,1}, \ldots, w_{0,K})$, and the $r \times r$ residual covariance matrix $\boldsymbol{V}$.

**Require:** Convergence threshold, tol $\geq 0$, and an upper limit on the number of iterations, $t_{\max}$.

$t \leftarrow 0$
$\delta \leftarrow \infty$
$\mathrm{ELBO}^{(0)} \leftarrow F(q; \boldsymbol{V}, \boldsymbol{w}_0)$
**while** $\delta >$ tol and $t < t_{\max}$ **do**
    $t \leftarrow t + 1$
    Compute expected residuals, $\bar{\boldsymbol{R}} \leftarrow \boldsymbol{Y} - \boldsymbol{X}\bar{\boldsymbol{B}}$
    Initialize $\boldsymbol{W}$ to a $p \times K$ matrix of zeros
    **for** $j$ in $1, \ldots, p$ **do**
        Remove variable $j$ from expected residuals, $\bar{\boldsymbol{R}}_j \leftarrow \bar{\boldsymbol{R}} + \boldsymbol{x}_j \bar{\boldsymbol{b}}_j^{\intercal}$
        $(w_{1,1}^{(j)}, \ldots, w_{1,K}^{(j)}, \boldsymbol{b}_{1,1}^{(j)}, \ldots, \boldsymbol{b}_{1,K}^{(j)}, \boldsymbol{S}_{1,1}^{(j)}, \ldots, \boldsymbol{S}_{1,K}^{(j)}) \leftarrow \text{BMSR-mix}(\boldsymbol{x}_j, \bar{\boldsymbol{R}}_j, \boldsymbol{V}, \mathscr{S}_0, \boldsymbol{w}_0)$ ▷ See (58).
        Compute posterior mean, $\bar{\boldsymbol{b}}_j = \sum_{k=1}^{K} w_{1,k}^{(j)} \boldsymbol{b}_{1,k}^{(j)}$
        Compute posterior covariance, $\boldsymbol{S}_j = \sum_{k=1}^{K} w_{1,k}^{(j)} [\boldsymbol{b}_{1,k}^{(j)} (\boldsymbol{b}_{1,k}^{(j)})^{\intercal} + \boldsymbol{S}_{1,k}^{(j)}] - \bar{\boldsymbol{b}}_j \bar{\boldsymbol{b}}_j^{\intercal}$
        Store $w_{1,1}^{(j)}, \ldots, w_{1,K}^{(j)}$ in the $j$th row of $\boldsymbol{W}$
        Include variable $j$ in expected residuals, $\bar{\boldsymbol{R}} \leftarrow \bar{\boldsymbol{R}}_j - \boldsymbol{x}_j \bar{\boldsymbol{b}}_j^{\intercal}$
    Update prior weights, $\boldsymbol{w}_0 \leftarrow$ UPDATE-WEIGHTS($\boldsymbol{W}$) ▷ See (14).
    Update residual covariance, $\boldsymbol{V} \leftarrow$ UPDATE-RESID-COV($\boldsymbol{X}, \bar{\boldsymbol{R}}, \boldsymbol{S}_1, \ldots, \boldsymbol{S}_p$) ▷ See (12).
    $\mathrm{ELBO}^{(t)} \leftarrow F(q; \boldsymbol{V}, \boldsymbol{w}_0)$
    $\delta \leftarrow \mathrm{ELBO}^{(t)} - \mathrm{ELBO}^{(t-1)}$
**return** $\bar{\boldsymbol{B}}$, $\boldsymbol{V}$, $\boldsymbol{w}_0$, $\mathrm{ELBO}^{(t)}$

$O(nr^3)$.

#### *D.1. Computing the posterior distribution of the regression coefficients*

Our aims are to (i) compute posterior estimates of the *mr.mash* unknowns and (ii) adapt the *mr.mash* priors to the data. Both (i) and (ii) are computationally challenging in multiple linear regression models, especially in a multivariate framework such as *mr.mash*. (With some exceptions such as ridge regression.) Therefore, some numerical approximations are required. The most widely used numerical approximations are based on Monte Carlo methods, most commonly Markov chain Monte Carlo (MCMC) and variants of MCMC (Clyde et al., 2011; Dellaportas et al., 2002; Erbe et al., 2012; George and McCulloch, 1993; Gianola et al., 2009; Hoggart et al., 2008; Guan and Stephens, 2011; Meuwissen et al., 2001; Moser et al., 2015; Perez and de los Campos, 2014; Zanella and Roberts, 2019; Zhou et al., 2013). While MCMC and other Monte Carlo methods have allowed for a proliferation of different models for prediction, Monte Carlo methods typically involve a high computational effort, and therefore may limit our ability to apply *mr.mash* to genetic data sets with many genetic variants and many phenotypes. Another approach is based on stochastic search methods (Benner et al., 2016; Bottolo and Richardson, 2010; Bottolo et al., 2011, 2013; Hans et al., 2007; Wallace et al., 2015). But stochastic search only focuses on identifying highest-probability configurations, and therefore these approaches are not well suited to predicting $\boldsymbol{Y}$ and adapting the priors.

An alternative approach to MCMC is variational inference (VI) (Blei et al., 2017; Jordan et al., 1999). In VI, we treat posterior inference as an optimization problem; the aim is to find a distri-

**Algorithm 3** *mr.mash* with missing data

**Require:** $n \times p$ data matrix $\boldsymbol{X}$ (does not need to be centered). Optionally, $\boldsymbol{X}$ may be "standardized"—that is, scaled so that each column has unit standard deviation—which simplifies downstream posterior computations (see Sec. I).

**Require:** Set of $K$ covariance matrices, $\mathscr{S}_0$.

**Require:** Initial estimates of the posterior mean coefficients, stored as a $p \times r$ matrix, $\bar{\boldsymbol{B}}$, and initial estimate of the intercept, $\bar{\boldsymbol{b}}_0$ (a vector of length $r$).

**Require:** Initial estimate of $\boldsymbol{Y}_{\text{full}}$, the $n \times r$ response matrix containing the observed values and initial estimates of missing values.

**Require:** Initial estimates of the prior mixture weights, $\boldsymbol{w}_0 = (w_{0,1}, \ldots, w_{0,K})$, and the $r \times r$ residual covariance matrix $\boldsymbol{V}$.

**while** convergence criterion is not met **do**
- Center columns of $\boldsymbol{X}$ and store result in $\tilde{\boldsymbol{X}}$
- Impute missing values as posterior mean estimates given $\boldsymbol{b}_0, \bar{\boldsymbol{B}}, \boldsymbol{V}$, and store observed and missing values in $\boldsymbol{Y}_{\text{full}}$ (see eqs. 81, 82).
- Center columns of $\boldsymbol{Y}_{\text{full}}$ and store result in $\tilde{\boldsymbol{Y}}$
- Run Algorithm 2 with $t_{\max} \leftarrow 1$, in which $\boldsymbol{X}, \boldsymbol{Y}$ are set to $\tilde{\boldsymbol{X}}, \tilde{\boldsymbol{Y}}$
- Update intercept, $\hat{\boldsymbol{b}}_0 \leftarrow \frac{1}{n}(\boldsymbol{Y}_{\text{full}} - \boldsymbol{X}\bar{\boldsymbol{B}})^\intercal \boldsymbol{1}_n$ (see eq. 78)

**return** $\hat{\boldsymbol{b}}_0$, $\bar{\boldsymbol{B}}$, $\boldsymbol{Y}_{\text{full}}$, $\boldsymbol{V}$, $\boldsymbol{w}_0$

bution, $q(\boldsymbol{B})$, within a prescribed family of distributions, $\mathcal{Q}$, that best approximates the *mr.mash* posterior, $p(\boldsymbol{B} \mid \boldsymbol{X}, \boldsymbol{Y}, \boldsymbol{V}, \boldsymbol{w}_0, \mathscr{S}_0)$. The closeness of the approximate distribution to the true posterior is measured by the Kullback-Leibler (K-L) divergence, so the VI optimization problem is

$$\begin{array}{ll} \text{minimize} & D_{\text{KL}}(q(\boldsymbol{B}) \,\|\, p(\boldsymbol{B} \mid \boldsymbol{X}, \boldsymbol{Y}, \boldsymbol{V}, \boldsymbol{w}_0)) \\ \text{subject to} & q \in \mathcal{Q}, \end{array} \tag{5}$$

where $D_{\text{KL}}(q \,\|\, p)$ denotes the K-L divergence between distributions $q$ and $p$. (In this expression and in the expressions below, we omit the dependence on $\mathscr{S}_0$ since it is a prespecified model parameter and does not change.)

Since the K-L divergence is itself intractable, we instead work with a different objective function that is easier to compute but yields the same solutions as (5). This alternative objective is the "Evidence Lower Bound" (ELBO) (Blei et al., 2017):

$$F(q; \boldsymbol{V}, \boldsymbol{w}_0) := \log p(\boldsymbol{Y} \mid \boldsymbol{V}, \boldsymbol{w}_0) - D_{\text{KL}}(q(\boldsymbol{B}) \,\|\, p(\boldsymbol{B} \mid \boldsymbol{X}, \boldsymbol{Y}, \boldsymbol{V}, \boldsymbol{w}_0)) \tag{6}$$

Since the marginal likelihood $p(\boldsymbol{Y} \mid \boldsymbol{V}, \boldsymbol{w}_0)$ does not depend on $q$, minimizing the K-L divergence with respect to $q$ is equivalent to maximizing the ELBO:

$$\underset{q \in Q}{\operatorname{argmin}}\, D_{\text{KL}}(q(\boldsymbol{B}) \,\|\, p(\boldsymbol{B} \mid \boldsymbol{X}, \boldsymbol{Y}, \boldsymbol{V}, \boldsymbol{w}_0)) = \underset{q \in Q}{\operatorname{argmax}}\, F(q; \boldsymbol{V}, \boldsymbol{w}_0) \tag{7}$$

The advantage of the ELBO, however, is that, by straightforward algebraic manipulations the ELBO can be rewritten as

$$F(q; \boldsymbol{V}, \boldsymbol{w}_0) = E_q[\log p(\boldsymbol{Y} \mid \boldsymbol{X}, \boldsymbol{B}, \boldsymbol{V}, \boldsymbol{w}_0)] + E_q\left[\log\left(\frac{p(\boldsymbol{B}; \boldsymbol{w}_0)}{q(\boldsymbol{B})}\right)\right], \tag{8}$$

which will lead to tractable computations for specific choices of $\mathcal{Q}$.

To make computations feasible in large data sets, we introduce the assumption that the $\boldsymbol{b}_j$'s are independent in the approximate posterior:

$$q(\boldsymbol{B}) = \prod_{j=1}^{p} q_j(\boldsymbol{b}_j) \tag{9}$$

In the VI literature, this assumption is referred to as the "mean-field" approximation. In practice, this independence assumption has the effect of selecting at most one variable with high probability when multiple variables are strongly correlated with each other (Carbonetto and Stephens, 2012). This can be a problem if variable selection is the main goal (Wang et al., 2020), but it is less of a concern when phenotype prediction is the main goal, which is precisely our goal here. Under this independence assumption, we reexpress the optimization problem as

$$\hat{q}_1, \ldots, \hat{q}_p := \underset{q_1,\cdots,q_p}{\operatorname{argmax}} \; F(q_1, \ldots, q_j; \boldsymbol{V}, \boldsymbol{w}_0). \tag{10}$$

Optimizing the ELBO over all $q_1, \ldots, q_p$ simultaneously is difficult. But optimizing a single factor $q_j$ is relatively straightforward because it reduces to computing analytic posterior quantities for a Bayesian multivariate simple regression model. We formalize this statement in the following proposition.

PROPOSITION 1. Let $\bar{\boldsymbol{b}}_j$ be the expected value of $\boldsymbol{b}_j$ with respect to the approximate posterior $q_j$, let $\bar{\boldsymbol{R}}_j = \boldsymbol{Y} - \sum_{j' \neq j} \boldsymbol{x}_{j'} \bar{\boldsymbol{b}}_{j'}^{\intercal}$ be the expected residuals ignoring the $j$th variable, and define a function

$$\text{BMSR-mix}(\boldsymbol{x}, \boldsymbol{Y}, \boldsymbol{V}, \mathscr{S}_0, \boldsymbol{w}_0) := (w_{1,1}, \ldots, w_{1,K}, \boldsymbol{b}_{1,1}, \ldots, \boldsymbol{b}_{1,K}, \boldsymbol{S}_{1,1}, \ldots, \boldsymbol{S}_{1,K})$$

that returns the posterior distribution of $\boldsymbol{b}$ under the Bayesian multivariate simple regression model with a mixture-of-normals prior (see Proposition 5 for the exact definition of BMSR-mix). Then we have that

$$\begin{aligned} \operatorname{argmax}_{q_j} \; F(q_1, \ldots, q_j; \boldsymbol{V}, \boldsymbol{w}_0) &= \text{BMSR-mix}(\boldsymbol{x}_j, \bar{\boldsymbol{R}}_j, \boldsymbol{V}, \mathscr{S}_0, \boldsymbol{w}_0) \\ &:= (w_{1,1}^{(j)}, \ldots, w_{1,K}^{(j)}, \boldsymbol{b}_{1,1}^{(j)}, \ldots, \boldsymbol{b}_{1,K}^{(j)}, \boldsymbol{S}_{1,1}^{(j)}, \ldots, \boldsymbol{S}_{1,K}^{(j)}). \end{aligned} \tag{11}$$

This suggests a *co-ordinate ascent algorithm* for maximizing the ELBO, in which we update the $q_j$'s sequentially. In the next section, we turn to the question of updating the other model parameters, $\boldsymbol{w}_0$ and $\boldsymbol{V}$.

#### *D.2. Estimating $\boldsymbol{V}$ and $\boldsymbol{w}_0$*

We estimate $\boldsymbol{w}_0$ and $\boldsymbol{V}$ by maximizing the ELBO over $\boldsymbol{w}_0$ and $\boldsymbol{V}$, with $q$ fixed. This procedure can be viewed as an "EM-like" algorithm in which updating $q$ is an approximate E-step, and updating $\boldsymbol{w}_0$ and $\boldsymbol{V}$ is an M-step (Neal and Hinton, 1998).

For the residual covariance matrix, $\boldsymbol{V}$, the update is

$$\text{UPDATE-RESID-COV}(\boldsymbol{X}, \bar{\boldsymbol{R}}, \boldsymbol{S}_1, \ldots, \boldsymbol{S}_p) := \operatorname{argmax}_{\boldsymbol{V}} F(q; \boldsymbol{V}, \boldsymbol{w}_0) = \text{ERSS}/n, \tag{12}$$

in which "ERSS" is the expected residual sum of squares,

$$\text{ERSS} := E_q[\boldsymbol{R}^{\intercal}\boldsymbol{R}] = \bar{\boldsymbol{R}}^{\intercal}\bar{\boldsymbol{R}} + \sum_{j=1}^{p} \boldsymbol{x}_j^{\intercal}\boldsymbol{x}_j\boldsymbol{S}_j, \tag{13}$$

where $\boldsymbol{R} = \boldsymbol{Y} - \boldsymbol{X}\boldsymbol{B}$ is the $n \times r$ matrix of residuals, $\bar{\boldsymbol{R}} := E_q[\boldsymbol{R}] = \boldsymbol{Y} - \boldsymbol{X}\bar{\boldsymbol{B}}$, and $\boldsymbol{S}_j = \text{Cov}_q(\boldsymbol{b}_j)$. See Sec. G for a more detailed derivation of these expressions.

To derive the update for the mixture weights, $\boldsymbol{w}_0$, we consider an augmented representation of the *mr.mash* model. Maximizing the ELBO for this augmented model, denoted as $\tilde{F}(q; \boldsymbol{V}, \boldsymbol{w}_0)$, with $q$ fixed, yields the following update:

$$\text{UPDATE-WEIGHTS}(\boldsymbol{W}) := \operatorname{argmax}_{\boldsymbol{w}_0} \tilde{F}(q; \boldsymbol{V}, \boldsymbol{w}_0) = (\hat{w}_{0,1}, \ldots, \hat{w}_{0,K}), \tag{14}$$

in which $\hat{w}_{0,k} := \sum_{j=1}^{p} w_{1,k}^{(j)}/p$, $w_{1,k}^{(j)}$ is defined in (11), and $\boldsymbol{W}$ is a $p \times K$ matrix in which $w_{1,k}^{(j)}$ is stored in row $j$, column $k$ of the matrix. See Sec. G for a more detailed derivation of this update.

### E. Details of the methods compared

This section describes how the methods were run in the simulations and the GTEx case study. Note that we ran all methods standardizing $\boldsymbol{X}$ and without standardizing $\boldsymbol{Y}$. The one exception was the *Elastic Net* implemented in the `glmnet` package; for *Elastic Net*, we standardized both $\boldsymbol{X}$ and $\boldsymbol{Y}$.

#### E.1. Elastic Net

The *Elastic Net* method is a univariate multiple linear regression method that typically produces sparse estimates of the regression coefficients through a combination of $\ell_1$ and $\ell_2$ penalties (Zou and Hastie, 2005). Since the *Elastic Net* does not model correlations among effects of different phenotypes, it is only expected to be competitive with the multivariate regression methods when there is little benefit to modeling correlations, such as when genetic variant effects are rarely shared across phenotypes, or when the effects are already accurately estimated in the individual regressions.

We used the fast algorithms implemented in version 4.1.1 of the R package `glmnet` (Friedman et al., 2010) to fit *Elastic Net* models separately to each of the $r$ responses. The penalty strength parameter $\lambda$ was chosen via cross-validation, separately for each of the $r$ responses. The mixing parameter $\alpha$, which controls the tradeoff between the Lasso penalty ($\ell_1$-norm) and the ridge penalty ($\ell_2$-norm), was set to 0.5, which was the same setting used in PrediXcan (Gamazon et al., 2015). (Since the ground-truth effects were sparse, $\alpha$ mainly served to avoid degeneracies caused by strong correlations between SNPs.) We used the coefficient estimates at the value of $\lambda$ minimizing the mean cross-validation error (`lambda.min`). We performed the cross-validation by calling `cv.glmnet` with `family = "gaussian"`, `alpha = 0.5`, and with all remaining arguments kept at their default values.

#### E.2. Group Lasso

The *Group Lasso* (Yuan and Lin, 2006) is a penalized multivariate regression method, similar in some respects to the *Elastic Net*, except that it uses a "$\ell_1/\ell_2$" penalty which gives greater preference to variables that have an effect in multiple phenotypes. As far as we know, the *Group Lasso* has not been used in the TWAS setting. However, we included *Group Lasso* in our comparisons because it has a very fast implementation in the R package `glmnet` (Friedman et al., 2010), and for our application we expect it to perform well in settings where effects are shared across many phenotypes. On the other hand, since the $\ell_1/\ell_2$ penalty does not specifically account for sharing of effect sizes, nor can it model sharing of effects among subsets of phenotypes, we expect that the gains in accuracy will be weaker in some settings. Also note that current implementations of the *Group Lasso* cannot handle missing data so we did not include the *Group Lasso* in our simulations with missing data.

We used the *Group Lasso* implemented in version 4.1.1 of the `glmnet` R package. We called `cv.glmnet` with `family = "mgaussian"`, `alpha = 1`, and with all remaining arguments kept at their default values. We used the coefficient estimates at the value of $\lambda$ minimizing the mean cross-validation error (`lambda.min`).

#### E.3. Sparse Multi-task Lasso

The *Sparse Multi-task Lasso* (Lee et al., 2010; Hu et al., 2019) is a variation on the *Group Lasso* with a more flexible penalty. The *Sparse Multi-task Lasso* was specifically developed for eQTL studies with missing data (Lee et al., 2010; Hu et al., 2019) and holds the promise of being able to adapt to a greater variety of data sets, and for these reasons it is a natural point of comparison to *mr.mash*. But in practice we found it was challenging to apply to the data sets considered in our simulations, and in particular in data sets without missing values it was much slower than the *Group Lasso* (implemented in the `glmnet` R package). Therefore, in practice the implementation of this method required compromises such as reducing the number of parameter settings in the cross-validation, and this may have reduced its performance.

The *Sparse Multi-task Lasso* minimizes the following penalized objective,

$$\ell^{\mathrm{smt}}(\boldsymbol{B}) = \frac{1}{2}\sum_{s=1}^{r} \|\boldsymbol{y}_s - \boldsymbol{X}\boldsymbol{b}_s\|_2^2 + \lambda_1 \sum_{s=1}^{r} \|\boldsymbol{b}_s\|_1 + \lambda_2 \sum_{j=1}^{p} \|\boldsymbol{b}_j\|_2, \tag{15}$$

where $\boldsymbol{y}_s$ denotes the $s$th column of $\boldsymbol{Y}$, $\boldsymbol{b}_s$ denotes the $s$th column of $\boldsymbol{B}$, and $\boldsymbol{b}_j$ denotes the $j$th row of $\boldsymbol{B}$. This formulation is equivalent to the objective function given in Hu et al. (2019) up to a scaling of the penalty. This was previously studied as the "sparse multi-task lasso problem" (Lee et al., 2010). The Lasso penalty on the *columns of* $\boldsymbol{B}$ encourages fewer predictors to be selected (Tibshirani, 1996), whereas the Group Lasso penalty on the *rows of* $\boldsymbol{B}$ encourages the same predictors to be chosen for different responses (Yuan and Lin, 2006).

To set the penalty parameters, we took a simple $k$-fold cross-validation approach. We used a uniformly spaced grid of candidate values on the log-scale for each of $\lambda_1$ and $\lambda_2$, evaluated the mean-squared error for each fold and each combination of $\lambda_1, \lambda_2$, and then selected the combination that had the smallest mean-squared error (averaged across all folds).

The implementation of the *Sparse Multi-task Lasso* made available in UTMOST (Hu et al., 2019) is difficult to use because of the nonstandard way it requires the inputs to be formatted. We also found that the memory usage could be very high when $\boldsymbol{Y}$ had missing values. The main UTMOST Python interface is available from `https://github.com/Joker-Jerome/UTMOST` and the core code implementing the *Sparse Multi-task Lasso* is available from `https://github.com/yiminghu/CTIMP`. Later, another implementation of the *Sparse Multi-task Lasso* was developed in Li et al. (2021) (see `https://github.com/RitchieLab/multi_tissue_twas_sim`). This newer implementation, while easier to use, does not allow for missing values in $\boldsymbol{Y}$. Due to these considerations, we developed our own Python implementation of the *Sparse Multi-task Lasso* (available at `https://github.com/aksarkar/mtlasso`). Our Python implementation is slower than previous implementations (S6 Fig), in part owing to the core computations not being implemented in C++. On the other hand, our implementation is easier to use, allows for missing values, and, in simulated data sets, performs similarly to the `multi_tissue_twas_sim` implementation (S5 Fig). For additional details on these implementations (which differ in the initialization method and the grid of tuning parameters, among other things) and simulations, see Morgante et al. (2023) (see in particular the file `mrmash_vs_mtlasso_vs_utmost.html`) and `https://users.rcc.uchicago.edu/~aksarkar/gtex-pred` (see in particular "Sparse multi-task lasso").

For $\boldsymbol{Y}$ with missing data, one complication is that the correct updates for $\boldsymbol{B}$ are expensive. Following the suggestion by Hu et al. (2019), the `mtlasso` implements an approximate update for each column $s = 1, \ldots, r$ of $\boldsymbol{B}$ in which $(\boldsymbol{X}^{(s)})^\intercal \boldsymbol{X}^{(s)}$ is replaced with $\boldsymbol{X}^\intercal \boldsymbol{X}$, in which $\boldsymbol{X}^{(s)}$ denotes the matrix containing the rows of $\boldsymbol{X}$ corresponding to non-missing entries of $\boldsymbol{y}_s$.

### E.4. mr.mash

In the simulations with full data, we obtained an initial estimate of $\bar{\boldsymbol{B}}$ by running the *Group Lasso* on the same data, then setting $\bar{\boldsymbol{B}}$ to the *Group Lasso* coefficient estimates. We then obtained an initial estimate of $\boldsymbol{V}$ as the sample covariance of the *Group Lasso* residuals. The prior mixture weight for the "spike", $w_{0,0}$ (see eq. 16), was initialized to the proportion of *Group Lasso* coefficients that were zero, and the remaining prior mixture components were assigned equal weights. In the *mr.mash* model fitting, we estimated both $\boldsymbol{w}_0$ and $\boldsymbol{V}$, but forced $\boldsymbol{V}$ to be diagonal. We ran the *mr.mash* model fitting algorithm with a convergence tolerance of 0.01.

For simulations with missing data, we ran *mr.mash* in the same way as in the simulations with full data, except for two differences. First, since the sample covariance of the residuals cannot be computed when there are missing values in $\boldsymbol{Y}$, we initialized $\boldsymbol{V}$ using `flashr` (Wang and Stephens, 2021), which can cope with missing values. Second, since the `glmnet` implementation of the *Group Lasso* does not allow missing values in $\boldsymbol{Y}$, we instead ran the *Elastic Net* separately on each column of $\boldsymbol{Y}$, and used the *Elastic Net* coefficient estimates to initialize $\bar{\boldsymbol{B}}$.

For the GTEx case study, we ran *mr.mash* in mostly the same way as in the simulations with missing data. Two differences were: (1) we allowed correlated residuals (*i.e.*, non-diagonal $\boldsymbol{V}$); and (2) to speed up the model fitting, at each outer-loop iteration beyond the initial 15 iterations we automatically dropped from the mixture prior any components with mixture weight smaller than $10^{-8}$.

#### E.4.1. Prior covariance matrices

We defined the covariance matrices as combinations of different scaling factors $\omega_l \geq 0$ and normalized covariance matrices, $\boldsymbol{U}_{0,t}$. (By "normalized", we mean that the largest diagonal element was always

1.) Therefore, the mixture prior was

$$\boldsymbol{b}_j \mid \boldsymbol{w}_0, \boldsymbol{\omega}, \mathscr{U}_0 \sim w_{0,0}\delta_{\mathbf{0}} + \sum_{l=1}^{L}\sum_{t=1}^{T} w_{0,l,t} N_r(\mathbf{0}, \omega_l^2 \boldsymbol{U}_{0,t}), \tag{16}$$

where $\delta_{\mathbf{0}}$ is the delta mass at zero (the "spike"). The scaling factors $\omega_1, \ldots, \omega_L$ were chosen so that they were spaced uniformly on the log-scale, with a grid spacing of $\sqrt{2}$. The steps we took to determine the end points of the grid are described in Urbut et al. (2019).

In the next two sections we describe how the normalized covariance matrices $\boldsymbol{U}_{0,1}, \ldots, \boldsymbol{U}_{0,T}$ were specified.

#### *E.4.2. Canonical covariance matrices*

When we ran *mr.mash* using "canonical" covariance matrices, the following matrices were included in the prior (16):

- The identity matrix, $\boldsymbol{I}_r$, modeling the case in which all effects are independent.
- The "equal effects" matrix, an $r \times r$ matrix of all ones, which models the case in which all effects are the same.
- Rank-1 matrices modeling tissue-specific effects. They are of the form $\boldsymbol{c}_s\boldsymbol{c}_s^{\intercal}$, in which $c_s$ is a vector of length $r$ containing all zeros except for a 1 in the $s$th position. There were $r$ of these matrices.
- Matrices with ones on the diagonal and $\sigma$ on the off-diagonal, where $\sigma$ is 0.25, 0.5 or 0.75. There were three three of these matrices.

In total, $r + 5$ canonical matrices were included in the prior. So for the simulations the prior included 15 canonical matrices.

#### *E.4.3. Data-driven matrices*

Data-driven matrices are, as the term implies, adapted to the data, in contrast to the canonical matrices which are the same for every data set. The idea is that these data-driven covariance matrices should capture the principal patterns of effect sharing patterns across the $r$ tissues. Estimating the data-driven covariance matrices involves combining information across many SNPs and many genes.

For the simulations, when data-driven matrices were included in the prior (16), we estimated the data-driven matrices following the analysis procedure largely as it was described in Urbut et al. (2019). In brief, we prepared the summary statistics (least-squares effect estimates and corresponding standard errors) using training sets (real genotypes $\boldsymbol{X}$ and simulated expression levels $\boldsymbol{Y}$) for all genes, then we used these summary statistics to estimate the data-driven covariance matrices using Extreme Deconvolution (ED) (Bovy et al., 2011).

We used ED to estimate data-driven covariances because it is an EM algorithm for fitting mixture models of the form (2) in which the multivariate effect vectors $\boldsymbol{b}_j$ are treated as unknown, and "noisy" estimates of the $\boldsymbol{b}_j$'s are observed. In particular, ED estimates both the mixture weights $w_{0,k}$ and the covariance matrices $\boldsymbol{S}_{0,k}$ in (2). Once the ED algorithm converged to a solution, we extracted the covariance matrices estimated by ED to use as the data-driven covariances $\boldsymbol{U}_{0,t}$ in the prior.

To run the ED algorithm, one needs to provide initial estimates of the covariance matrices. We obtained rough initial estimates by running PCA and the flexible factor analysis methods implemented in the R package `flashr` (Wang and Stephens, 2021), as well as one covariance estimate initialized to the empirical covariance matrix (that is, the maximum-likelihood estimate of the covariance). For the PCA-based initial estimates, we simply computed a singular value decomposition (implemented by R function `svd`) of a matrix formed by selected least-squares estimates of the effects $\boldsymbol{b}_j$, then obtained 3 rank-1 matrices from the top 3 PCs, and a rank-3 matrix from the linear combination of the top 3 PCs (so 4 PCA-based covariance matrices in total). After running `flashr`, we formed rank-1 covariance matrices from the estimated factors. Since `flashr` adaptively determines the number of factors—essentially, `flashr` will remove factors during the back-fitting stage if setting the factor to zero does not make the model fit worse—the number of covariances fit by ED was different for each data set. By automatically determining the number of factors, the idea is that the number of factors (and therefore the number of components in the mixture prior) should approximately reflect the complexity the effect sharing across tissues. Note that for the Equal Effects and Equal Effects + Null simulations, the final result was always a mixture model with 9 mixture components (*i.e.*, 9 covariance matrices, 4 initialized with PCA, 4 initialized with `flashr`, and 1 initialized to the empirical covariance matrix).

The code for preprocessing the summary statistics is available from `prepare_sumstats_for_ED_prior.R` in Morgante et al. (2023), and the code implementing the remaining steps of this analysis is available at `https://github.com/stephenslab/gtexresults/blob/master/workflows/mashr_flashr_workflow.ipynb` (we used the "prior" workflow, version 2570aa6 from 2019-10-03). The one difference between the analysis of Urbut et al. (2019) and ours is that we replaced the sparse factor analysis (SFA) method (Engelhardt and Stephens, 2010) with `flashr` (Wang and Stephens, 2021).

For the GTEx application, we made a couple small improvements to this pipeline. The key difference was the use of in-development model fitting algorithms that improve on the EM algorithms used in Extreme Deconvolution (Bovy et al., 2011), and are implemented in the `udr` package available at `https://github.com/stephenslab/udr`. The script implementing this modified pipeline is available at `https://github.com/cumc/bioworkflows/blob/master/multivariate-fine-mapping/mixture_prior.ipynb` (we used the "extract_effects" workflow followed by the "ud" workflowr in the script version with git commit id 5b7c688, with flags `--ud-method ted --ud-tol-lik 1e-3 --mixture-components flash flash_nonneg pca`). Since there were 5 training sets in the GTEx application, we ran this pipeline 5 times. In each training set, the pipeline produced a mixture model with 33–35 components.

### F. Posterior computations for multivariate simple regression models

Here we derive elementary posterior expressions that will be used later on to develop the posterior computations for *mr.mash*.

#### F.1. Multivariate simple regression

We begin by deriving posterior computations for a multivariate regression with a single explanatory variable. Initially, we do not include an intercept.

The model is

$$\begin{aligned} \boldsymbol{Y} &= \boldsymbol{x}\boldsymbol{b}^\intercal + \boldsymbol{E} \\ \boldsymbol{E} &\sim MN_{n\times r}(\boldsymbol{0}, \boldsymbol{I}_n, \boldsymbol{V}), \end{aligned} \tag{17}$$

or more concisely

$$\boldsymbol{Y} \sim MN_{n\times r}(\boldsymbol{x}\boldsymbol{b}^\intercal, \boldsymbol{I}_n, \boldsymbol{V}), \tag{18}$$

where $\boldsymbol{Y} \in \mathbb{R}^{n\times r}$ is a matrix of observed responses for $n$ samples across $r$ conditions, $\boldsymbol{x}$ is a vector of $n$ observations for a single explanatory variable, $\boldsymbol{b} \in \mathbb{R}^r$ is the (unknown) vector of regression for the $r$ conditions, $\boldsymbol{V} \in \mathbb{S}^r_{++}$ is an invertible covariance matrix, and $MN_{n\times r}(\boldsymbol{M}, \boldsymbol{U}, \boldsymbol{V})$ is the matrix normal distribution (Dawid, 1981; Gupta and Nagar, 2000) with mean $\boldsymbol{M}$ and covariance matrices $\boldsymbol{U}$ and $\boldsymbol{V}$ (these are matrices of dimension $n \times r$, $n \times n$ and $r \times r$, respectively).

In the following, we give the expression for the matrix-normal likelihood and relate it to the more familiar multivariate normal distribution. The likelihood for the multivariate regression model (18) is

$$\begin{aligned} \ell(\boldsymbol{b}; \boldsymbol{x}, \boldsymbol{Y}, \boldsymbol{V}) &:= p(\boldsymbol{Y} \mid \boldsymbol{x}, \boldsymbol{b}, \boldsymbol{V}) \\ &= MN_{n\times r}(\boldsymbol{Y}; \boldsymbol{x}\boldsymbol{b}^\intercal, \boldsymbol{I}_n, \boldsymbol{V}) \\ &= |2\pi\boldsymbol{V}|^{-n/2} \exp\{-\tfrac{1}{2}\mathrm{tr}[\boldsymbol{V}^{-1}(\boldsymbol{Y} - \boldsymbol{x}\boldsymbol{b}^\intercal)^\intercal(\boldsymbol{Y} - \boldsymbol{x}\boldsymbol{b}^\intercal)]\}. \end{aligned} \tag{19}$$

Given $\boldsymbol{V}$, the least-squares estimate of $\boldsymbol{b}$, denoted $\hat{\boldsymbol{b}}$, which is also the value of $\boldsymbol{b}$ maximizing the likelihood (19), and its covariance, $\hat{\boldsymbol{S}}$, are

$$\hat{\boldsymbol{b}} := \hat{\boldsymbol{b}}(\boldsymbol{x}, \boldsymbol{Y}, \boldsymbol{V}) = \tfrac{\boldsymbol{Y}^\intercal \boldsymbol{x}}{\boldsymbol{x}^\intercal \boldsymbol{x}} \tag{20}$$

$$\hat{\boldsymbol{S}} := \hat{\boldsymbol{S}}(\boldsymbol{x}, \boldsymbol{Y}, \boldsymbol{V}) = \tfrac{\boldsymbol{V}}{\boldsymbol{x}^\intercal \boldsymbol{x}}. \tag{21}$$

Using these quantities, the likelihood (19) can be rewritten as

$$\ell(\boldsymbol{b}; \boldsymbol{x}, \boldsymbol{Y}, \boldsymbol{V}) = |2\pi \boldsymbol{V}|^{-n/2} \exp\{-\tfrac{1}{2}[\mathrm{tr}(\boldsymbol{Y}\boldsymbol{V}^{-1}\boldsymbol{Y}^\intercal) + (\boldsymbol{b} - \hat{\boldsymbol{b}})^\intercal \hat{\boldsymbol{S}}^{-1}(\boldsymbol{b} - \hat{\boldsymbol{b}}) - \hat{\boldsymbol{b}}^\intercal \hat{\boldsymbol{S}}^{-1}\hat{\boldsymbol{b}}]\}. \tag{22}$$

This expression is proportional to a multivariate normal density on $\boldsymbol{b}$, and in particular we have that

$$\ell(\boldsymbol{b}; \boldsymbol{x}, \boldsymbol{Y}, \boldsymbol{V}) \propto N_r(\boldsymbol{b}; \hat{\boldsymbol{b}}, \hat{\boldsymbol{S}}), \tag{23}$$

where $N_n(\boldsymbol{\theta}; \boldsymbol{\mu}, \boldsymbol{\Sigma})$ denotes the multivariate normal density at $\boldsymbol{\theta} \in \mathbb{R}^n$ with mean $\boldsymbol{\mu}$ and covariance $\boldsymbol{\Sigma}$.

#### *F.2. Bayesian multivariate simple regression with a normal prior*

In the following proposition, we apply the above results for the multivariate simple regression to a Bayesian multivariate simple regression model with a normal prior.

PROPOSITION 2 (BAYESIAN MULTIVARIATE SIMPLE REGRESSION WITH A NORMAL PRIOR). *Consider the multivariate simple regression model* (18) *with a multivariate normal prior on the regression coefficients,*

$$\boldsymbol{b} \mid \boldsymbol{S}_0 \sim N_r(\boldsymbol{0}, \boldsymbol{S}_0), \tag{24}$$

*where* $\boldsymbol{S}_0 \in \mathbb{S}_+^r$ *is a (possibly singular) covariance matrix. The posterior of* $\boldsymbol{b}$ *is*

$$\boldsymbol{b} \mid \boldsymbol{x}, \boldsymbol{Y}, \boldsymbol{V}, \boldsymbol{S}_0 \sim N_r(\boldsymbol{b}_1, \boldsymbol{S}_1), \tag{25}$$

*where*

$$\boldsymbol{b}_1 := \boldsymbol{b}_1(\boldsymbol{x}, \boldsymbol{Y}, \boldsymbol{V}, \boldsymbol{S}_0) = \boldsymbol{S}_1 \hat{\boldsymbol{S}}^{-1} \hat{\boldsymbol{b}} \tag{26}$$

$$\boldsymbol{S}_1 := \boldsymbol{S}_1(\boldsymbol{x}, \boldsymbol{Y}, \boldsymbol{V}, \boldsymbol{S}_0) = (\boldsymbol{S}_0^{-1} + \hat{\boldsymbol{S}}^{-1})^{-1}. \tag{27}$$

*The Bayes Factor (BF) comparing this model against the null model* ($\boldsymbol{b} = \boldsymbol{0}$)*, which is defined as the ratio of the two likelihoods,*

$$\begin{aligned}\mathrm{BF}(\boldsymbol{x}, \boldsymbol{Y}, \boldsymbol{V}, \boldsymbol{S}_0) &= \frac{p(\boldsymbol{Y} \mid \boldsymbol{x}, \boldsymbol{V}, \boldsymbol{S}_0)}{p(\boldsymbol{Y} \mid \boldsymbol{x}, \boldsymbol{b} = \boldsymbol{0}, \boldsymbol{V})} \\ &= \frac{\int \ell(\boldsymbol{b}; \boldsymbol{x}, \boldsymbol{Y}, \boldsymbol{V})\, p(\boldsymbol{b} \mid \boldsymbol{S}_0)\, d\boldsymbol{b}}{p(\boldsymbol{Y} \mid \boldsymbol{x}, \boldsymbol{b} = \boldsymbol{0}, \boldsymbol{V})},\end{aligned} \tag{28}$$

*works out to*

$$\mathrm{BF}(\boldsymbol{x}, \boldsymbol{Y}, \boldsymbol{V}, \boldsymbol{S}_0) = \frac{|\hat{\boldsymbol{S}}|^{1/2}}{|\boldsymbol{S}_0 + \hat{\boldsymbol{S}}|^{1/2}} \exp\{\tfrac{1}{2} \hat{\boldsymbol{b}}^\intercal \hat{\boldsymbol{S}}^{-1} \boldsymbol{S}_1 \hat{\boldsymbol{S}}^{-1} \hat{\boldsymbol{b}}\} \tag{29}$$

$$= \frac{|\hat{\boldsymbol{S}}|^{1/2}}{|\boldsymbol{S}_0 + \hat{\boldsymbol{S}}|^{1/2}} \exp\{\tfrac{1}{2} \boldsymbol{b}_1^\intercal \boldsymbol{S}_1^{-1} \boldsymbol{b}_1\}. \tag{30}$$

*The same BF can also be equivalently expressed as a ratio of two multivariate normal densities,*

$$\mathrm{BF}(\boldsymbol{x}, \boldsymbol{Y}, \boldsymbol{V}, \boldsymbol{S}_0) = \frac{N_r(\hat{\boldsymbol{b}}; \boldsymbol{0}, \boldsymbol{S}_0 + \hat{\boldsymbol{S}})}{N_r(\hat{\boldsymbol{b}}; \boldsymbol{0}, \hat{\boldsymbol{S}})}. \tag{31}$$

#### F.3. Multivariate simple regression with an intercept

Now we extend the multivariate simple regression model (18) to include an intercept. We show that including the intercept in the model is equivalent to "centering" $\boldsymbol{x}$ and the columns of $\boldsymbol{Y}$ so that they all have means of zero. This equivalence can be understood from two different perspectives: from a point-estimation perspective, centering $\boldsymbol{x}$ and the columns of $\boldsymbol{Y}$ is equivalent to computing a maximum-likelihood estimate of the intercept; from a Bayesian averaging point-of-view, it is equivalent to integrating out the intercept with respect to an (improper) uniform prior (see also George and McCulloch 1997). These results are summarized in Proposition 3.

The multivariate simple regression model with an intercept is

$$\boldsymbol{Y} \sim MN_{n\times r}(\mathbf{1}\boldsymbol{\mu}^\intercal + \boldsymbol{x}\boldsymbol{b}^\intercal, \boldsymbol{I}_n, \boldsymbol{V}), \tag{32}$$

in which $\mathbf{1}$ is a vector of ones of length $n$, and $\boldsymbol{\mu} \in \mathbb{R}^r$ is the (unknown) intercept. The likelihood of $\boldsymbol{\mu}, \boldsymbol{b}$ under this model is

$$\begin{aligned}\ell(\boldsymbol{\mu}, \boldsymbol{b}; \boldsymbol{x}, \boldsymbol{Y}, \boldsymbol{V}) &:= p(\boldsymbol{Y} \mid \boldsymbol{x}, \boldsymbol{\mu}, \boldsymbol{b}, \boldsymbol{V}) \\ &= |2\pi\boldsymbol{V}|^{-n/2}\exp\{-\tfrac{1}{2}\mathrm{tr}[\boldsymbol{V}^{-1}(\boldsymbol{Y} - \mathbf{1}\boldsymbol{\mu}^\intercal - \boldsymbol{x}\boldsymbol{b}^\intercal)^\intercal(\boldsymbol{Y} - \mathbf{1}\boldsymbol{\mu}^\intercal - \boldsymbol{x}\boldsymbol{b}^\intercal)]\}. \end{aligned}\tag{33}$$

Proposition 3 (Multivariate simple regression with an intercept). *Consider the multivariate simple regression* (32). *The least-squares estimate of $\boldsymbol{\mu}$—which is also the value of $\boldsymbol{\mu}$ maximizing the likelihood $\ell(\boldsymbol{\mu}, \boldsymbol{b}; \boldsymbol{x}, \boldsymbol{Y}, \boldsymbol{V})$—and its covariance matrix are*

$$\hat{\boldsymbol{\mu}} := \hat{\boldsymbol{\mu}}(\boldsymbol{x}, \boldsymbol{Y}, \boldsymbol{V}, \boldsymbol{b}) = \bar{\boldsymbol{y}} - \bar{x}\boldsymbol{b} \tag{34}$$

$$\hat{\boldsymbol{S}}_\mu := \hat{\boldsymbol{S}}_\mu(\boldsymbol{x}, \boldsymbol{Y}, \boldsymbol{V}, \boldsymbol{b}) = \tfrac{1}{n}\boldsymbol{V}, \tag{35}$$

*in which $\bar{x} = \frac{1}{n}\boldsymbol{x}^\intercal\mathbf{1} = \frac{1}{n}\sum_{i=1}^n x_i$ is the sample mean of $\boldsymbol{x}$, and $\bar{\boldsymbol{y}} = \frac{1}{n}\boldsymbol{Y}^\intercal\mathbf{1}$ is the vector containing the column means of $\boldsymbol{Y}$.*

*The profile likelihood for $\boldsymbol{b}$ is*

$$\begin{aligned}\ell^*(\boldsymbol{b}; \boldsymbol{x}, \boldsymbol{Y}, \boldsymbol{V}) &:= \max_{\boldsymbol{\mu}} \ell(\boldsymbol{\mu}, \boldsymbol{b}; \boldsymbol{x}, \boldsymbol{Y}, \boldsymbol{V}) \\ &= \ell(\boldsymbol{b}; \tilde{\boldsymbol{x}}, \tilde{\boldsymbol{Y}}, \boldsymbol{V}),\end{aligned} \tag{36}$$

*in which $\tilde{\boldsymbol{x}} := \boldsymbol{x} - \bar{x}\mathbf{1}$ and $\tilde{\boldsymbol{Y}} := \boldsymbol{Y} - \mathbf{1}\bar{\boldsymbol{y}}^\intercal$ are the centered $\boldsymbol{x}$ and $\boldsymbol{Y}$. In other words, the profile likelihood for the multivariate simple regression with an intercept is the same as the likelihood for the multivariate simple regression* without *an intercept if we first center $\boldsymbol{x}$ and $\boldsymbol{Y}$. Centering $\boldsymbol{x}$ and $\boldsymbol{Y}$ is therefore equivalent to including an intercept in the multivariate regression and estimating the intercept by maximum-likelihood.*

*Next, consider Bayesian calculations for $\boldsymbol{\mu}$ with a multivariate normal prior, $\boldsymbol{\mu} \mid \boldsymbol{S}_{0\mu} \sim N_r(0, \boldsymbol{S}_{0\mu})$, in which $\boldsymbol{S}_{0\mu} \in \mathbb{S}_+^r$ is a (possibly singular) covariance matrix. The posterior for $\boldsymbol{\mu}$ conditioned on $\boldsymbol{b}$ is*

$$\boldsymbol{\mu} \mid \boldsymbol{x}, \boldsymbol{Y}, \boldsymbol{V}, \boldsymbol{S}_{0\mu}, \boldsymbol{b} \sim N_r(\boldsymbol{\mu}_1, \boldsymbol{S}_{1\mu}), \tag{37}$$

*where*

$$\boldsymbol{\mu}_1 := \boldsymbol{\mu}_1(\boldsymbol{x}, \boldsymbol{Y}, \boldsymbol{V}, \boldsymbol{S}_{0\mu}, \boldsymbol{b}) = \boldsymbol{S}_{1\mu}\hat{\boldsymbol{S}}_\mu^{-1}\hat{\boldsymbol{\mu}} \tag{38}$$

$$\boldsymbol{S}_{1\mu} := \boldsymbol{S}_{1\mu}(\boldsymbol{x}, \boldsymbol{Y}, \boldsymbol{V}, \boldsymbol{S}_{0\mu}, \boldsymbol{b}) = (\boldsymbol{S}_{0\mu}^{-1} + \hat{\boldsymbol{S}}_\mu^{-1})^{-1}. \tag{39}$$

*The marginal likelihood obtained by averaging over the intercept is*

$$\begin{aligned}\ell^\star(\boldsymbol{b}; \boldsymbol{x}, \boldsymbol{Y}, \boldsymbol{V}, \boldsymbol{S}_{0\mu}) &:= \textstyle\int \ell(\boldsymbol{\mu}, \boldsymbol{b}; \boldsymbol{x}, \boldsymbol{Y}, \boldsymbol{V})\, p(\boldsymbol{\mu} \mid \boldsymbol{S}_{0\mu})\, d\boldsymbol{\mu} \\ &= |2\pi\boldsymbol{V}|^{-n/2}|\boldsymbol{S}_{0\mu}^{-1}\boldsymbol{S}_{1\mu}|^{1/2}\exp\{\tfrac{1}{2}\boldsymbol{\mu}_1^\intercal\boldsymbol{S}_{1\mu}^{-1}\boldsymbol{\mu}_1 - \tfrac{1}{2}\mathrm{tr}[\boldsymbol{V}^{-1}(\boldsymbol{Y} - \boldsymbol{x}\boldsymbol{b}^\intercal)^\intercal(\boldsymbol{Y} - \boldsymbol{x}\boldsymbol{b}^\intercal)]\}.\end{aligned} \tag{40}$$

*In the special case of an (improper) uniform prior on* $\boldsymbol{\mu}$, *defined as* $\boldsymbol{\mu} \sim N_r(0, \boldsymbol{S}_{0\mu})$ *with* $\boldsymbol{S}_{0\mu}^{-1} \to 0$, *the posterior mean reduces to the least-squares estimate* $\boldsymbol{\mu}_1 = \hat{\boldsymbol{\mu}}$, *and* $\boldsymbol{S}_{1\mu} = \hat{\boldsymbol{S}}_{\mu}$, *and the marginal likelihood simplifies to*

$$\begin{aligned}\ell^{\star}(\boldsymbol{b}; \boldsymbol{x}, \boldsymbol{Y}, \boldsymbol{V}, \boldsymbol{S}_{0\mu}) &= |2\pi \boldsymbol{V}|^{-n/2} |\boldsymbol{S}_{0\mu}^{-1} \hat{\boldsymbol{S}}_{\mu}|^{1/2} \exp\left\{\tfrac{1}{2}\hat{\boldsymbol{\mu}}^{\intercal} \hat{\boldsymbol{S}}_{\mu}^{-1} \hat{\boldsymbol{\mu}} - \tfrac{1}{2}\mathrm{tr}\big[\boldsymbol{V}^{-1}(\boldsymbol{Y} - \boldsymbol{x}\boldsymbol{b}^{\intercal})^{\intercal}(\boldsymbol{Y} - \boldsymbol{x}\boldsymbol{b}^{\intercal})\big]\right\} \\ &= |\boldsymbol{S}_{0\mu}^{-1} \hat{\boldsymbol{S}}_{\mu}|^{1/2} \times \ell(\boldsymbol{b}; \tilde{\boldsymbol{x}}, \tilde{\boldsymbol{Y}}, \boldsymbol{V}). \end{aligned} \tag{41}$$

*In other words, the marginal likelihood for multivariate regression with an intercept when we use an improper uniform prior for the intercept is the same (up to a constant of proportionality) as the likelihood for multivariate regression* without *an intercept when we first center* $\boldsymbol{x}$ *and* $\boldsymbol{Y}$.

A proof of Proposition 3 is given in Sec. J.

#### F.4. Bayesian multivariate simple regression with a mixture prior

Here we extend the Bayesian multivariate simple regression with a normal prior to the multivariate simple regression with a mixture-of-normals prior,

$$\boldsymbol{b} \mid \mathscr{S}_0, \boldsymbol{w}_0 \sim \sum_{k=1}^{K} w_{0,k} N_r(\boldsymbol{0}, \boldsymbol{S}_{0,k}), \tag{42}$$

in which $\mathscr{S}_0 := \{\boldsymbol{S}_{0,1}, \ldots, \boldsymbol{S}_{0,K}\}$, each $\boldsymbol{S}_{0,k} \in \mathbb{S}_{+}^{r}$ is a (possibly singular) covariance matrix, for $k = 1, \ldots, K$, and the mixture weights $\boldsymbol{w}_0 = (w_{0,1}, \ldots, w_{0,K})$ are non-negative and sum to 1. The normal prior is a special case of (42) when $K = 1$.

To facilitate derivation of the posterior computations, we introduce the following data augmentation that recovers (42) after averaging, or integrating, over the latent variable $\gamma$:

$$\begin{aligned} p(\gamma = k \mid \boldsymbol{w}_0) &= w_{0,k} \\ \boldsymbol{b} \mid \mathscr{S}_0, \gamma = k &\sim N_r(\boldsymbol{0}, \boldsymbol{S}_{0,k}). \end{aligned} \tag{43}$$

This augmented model allows us to conveniently reuse the posterior computations for the simpler models; in particular, posterior computations conditioned on $\gamma$ reduce to computations for the Bayesian multivariate regression model with a normal prior, as we show in the proposition below.

PROPOSITION 4. *Given* $\boldsymbol{w}_0$ *and* $\mathscr{S}_0$, *the Bayes factor comparing this model against the null model* $(\boldsymbol{b} = \boldsymbol{0})$ *is*

$$\begin{aligned} \mathrm{BF}^{\mathsf{mix}}(\boldsymbol{x}, \boldsymbol{Y}, \boldsymbol{V}, \mathscr{S}_0, \boldsymbol{w}_0) &= \frac{p(\boldsymbol{Y} \mid \boldsymbol{x}, \boldsymbol{V}, \mathscr{S}_0, \boldsymbol{w}_0)}{p(\boldsymbol{Y} \mid \boldsymbol{x}, \boldsymbol{b} = \boldsymbol{0}, \boldsymbol{V})} \\ &= \frac{\sum_{k=1}^{K} w_{0,k} N_r(\hat{\boldsymbol{b}}; \boldsymbol{0}, \boldsymbol{S}_{0,k} + \hat{\boldsymbol{S}})}{N_r(\hat{\boldsymbol{b}}; \boldsymbol{0}, \hat{\boldsymbol{S}})}. \end{aligned} \tag{44}$$

*The posterior mean and covariance of* $\boldsymbol{b}$ *are*

$$\boldsymbol{b}_1^{\mathsf{mix}} := \boldsymbol{b}_1^{\mathsf{mix}}(\boldsymbol{x}, \boldsymbol{Y}, \boldsymbol{V}, \mathscr{S}_0, \boldsymbol{w}_0) = \textstyle\sum_{k=1}^{K} w_{1,k} \boldsymbol{b}_{1,k} \tag{45}$$

$$\boldsymbol{S}_1^{\mathsf{mix}} := \boldsymbol{S}_1^{\mathsf{mix}}(\boldsymbol{x}, \boldsymbol{Y}, \boldsymbol{V}, \mathscr{S}_0, \boldsymbol{w}_0) = \textstyle\sum_{k=1}^{K} w_{1,k} (\boldsymbol{b}_{1,k} \boldsymbol{b}_{1,k}^{\intercal} + \boldsymbol{S}_{1,k}) - \boldsymbol{b}_1^{\mathsf{mix}} (\boldsymbol{b}_1^{\mathsf{mix}})^{\intercal}, \tag{46}$$

*in which the posterior assignment probabilities* $w_{1,k} := p(\gamma = k \mid \boldsymbol{x}, \boldsymbol{Y}, \boldsymbol{V}, \mathscr{S}_0, \boldsymbol{w}_0)$ *are given by*

$$w_{1,k} = \frac{w_{0,k} \mathrm{BF}(\boldsymbol{x}, \boldsymbol{Y}, \boldsymbol{V}, \boldsymbol{S}_{0,k})}{\sum_{k'=1}^{K} w_{0,k'} \mathrm{BF}(\boldsymbol{x}, \boldsymbol{Y}, \boldsymbol{V}, \boldsymbol{S}_{0,k'})}. \tag{47}$$

*Here we use* $\boldsymbol{b}_{1,k}$ *and* $\boldsymbol{S}_{1,k}$ *to denote the posterior mean and covariance of* $\boldsymbol{b}$ *given* $\gamma = k$; *they are given by* (26) *and* (27), *respectively, replacing* $\boldsymbol{S}_0$ *with* $\boldsymbol{S}_{0,k}$.

### G. Derivation of *mr.mash* variational algorithms with fully observed $Y$

To develop the *mr.mash* algorithm, we formulate an optimization problem, then we design an iterative procedure for solving the optimization problem. Here we assume that $\boldsymbol{Y}$ is fully observed; in Sec. H we extend these methods to allow for missing values in $\boldsymbol{Y}$. For convenience, we restate the *mr.mash* model for fully observed $\boldsymbol{Y}$:

$$\begin{aligned} &\boldsymbol{Y} \sim MN_{n\times r}(\boldsymbol{XB}, \boldsymbol{I}_n, \boldsymbol{V}) \\ &\boldsymbol{b}_j \overset{i.i.d.}{\sim} g, \text{ for } j = 1, \ldots, p, \end{aligned} \tag{48}$$

where $\boldsymbol{Y}$ is an $n \times r$ matrix containing $n$ observations of $r$ outcomes, $\boldsymbol{X}$ is an $n \times p$ matrix containing $n$ observations of $p$ explanatory variables, and $\boldsymbol{E} \in \mathbb{R}^{n\times r}$ is a matrix of residuals. The unknowns are the regression coefficients $\boldsymbol{B} \in \mathbb{R}^{p\times r}$, the residual covariance matrix $\boldsymbol{V} \in \mathbb{S}_+^r$, and the prior on the regression coefficients, $g$. We denote the regression coefficients for the $j$th explanatory variable by $\boldsymbol{b}_j = (b_{1j}, \ldots, b_{rj})^\intercal$.

We develop an empirical Bayes method for *mr.mash* in which the residual covariance $\boldsymbol{V}$ and prior $g$ are estimated, and a posterior distribution of $\boldsymbol{B}$ is computed conditioned on the estimates of $\boldsymbol{V}$ and $g$. This is formalized as the following optimization problem:

$$\begin{aligned} &\text{maximize} && F(q, g, \boldsymbol{V}; \boldsymbol{X}, \boldsymbol{Y}) \\ &\text{subject to} && g \in \mathcal{G}, q \in \mathcal{Q}, \boldsymbol{V} \in \mathbb{S}_+^r, \end{aligned} \tag{49}$$

where $\mathcal{G}$ is a prespecified set of prior distributions on $\boldsymbol{b}$, $\mathcal{Q}$ is a prespecified set of posterior distributions for $\boldsymbol{B}$ (possibly the set of *all* valid posterior distributions on $\mathbb{R}^r$), and $F(q, g, \boldsymbol{V}; \boldsymbol{X}, \boldsymbol{Y})$ is the "evidence lower bound" (ELBO), a lower bound to the marginal likelihood (Blei et al., 2017):

$$\begin{aligned} \log p(\boldsymbol{Y} \mid \boldsymbol{X}, \boldsymbol{V}, g) &= \log \textstyle\int p(\boldsymbol{Y} \mid \boldsymbol{X}, \boldsymbol{V}, \boldsymbol{B}) \prod_{j=1}^p g(\boldsymbol{b}_j)\, d\boldsymbol{B} \\ &\geq F(q, g, \boldsymbol{V}; \boldsymbol{X}, \boldsymbol{Y}) \\ &= E_q[\log p(\boldsymbol{Y} \mid \boldsymbol{X}, \boldsymbol{V}, \boldsymbol{B})] - \textstyle\sum_{j=1}^p D_{\mathrm{KL}}(q_j(\boldsymbol{b}_j) \,\|\, g(\boldsymbol{b}_j)), \end{aligned} \tag{50}$$

in which $D_{\mathrm{KL}}(f(\boldsymbol{x}) \,\|\, g(\boldsymbol{x})) := \int f(\boldsymbol{x}) \log\{f(\boldsymbol{x})/g(\boldsymbol{x})\}\, d\boldsymbol{x}$ is the Kullback-Leibler (K-L) divergence measure between probability distributions $f(\boldsymbol{x})$ and $g(\boldsymbol{x})$ on $\boldsymbol{x} \in \mathbb{R}^d$, and $q_j(\boldsymbol{b}_j)$ is the marginal posterior probability for $\boldsymbol{b}_j$. When $\mathcal{Q}$ is the set of all valid posterior distributions on $\mathbb{R}^{p\times r}$, the variational distribution $q \in \mathcal{Q}$ maximizing the ELBO (50), $\hat{q} := \mathrm{argmax}_{q \in \mathcal{Q}}\, F(q, g, \boldsymbol{V}; \boldsymbol{X}, \boldsymbol{Y})$, is the true posterior, $\hat{q}(\boldsymbol{B}) \propto p(\boldsymbol{Y} \mid \boldsymbol{X}, \boldsymbol{V}, \boldsymbol{B})\, g(\boldsymbol{B})$, and the ELBO at $\hat{q}$, $F(\hat{q}, g, \boldsymbol{V}; \boldsymbol{X}, \boldsymbol{Y})$, is equal to the marginal likelihood $\log p(\boldsymbol{Y} \mid \boldsymbol{X}, \boldsymbol{V}, g)$ (Jordan et al., 1999; Neal and Hinton, 1998; Wainwright and Jordan, 2007).

#### G.1. Estimating $V$

To estimate $\boldsymbol{V}$, we focus on the parts of the ELBO that involve $\boldsymbol{V}$:

$$F(q, g, \boldsymbol{V}; \boldsymbol{X}, \boldsymbol{Y}) = E_q[\log p(\boldsymbol{Y} \mid \boldsymbol{X}, \boldsymbol{V}, \boldsymbol{B})] + \sum_{j=1}^p E_q[\log g(\boldsymbol{b}_j)] + \text{const}, \tag{51}$$

where "const" is a placeholder for additional terms that do not involve $g$ or $\boldsymbol{V}$. This is the familiar expected complete log-likelihood that appears in derivations of EM algorithms (Neal and Hinton, 1998), the one difference being that the expectations are computed with respect to $q(\boldsymbol{B})$ instead of the exact posterior, $p(\boldsymbol{B} \mid \boldsymbol{X}, \boldsymbol{Y}, \boldsymbol{V}, g) \propto p(\boldsymbol{Y} \mid \boldsymbol{X}, \boldsymbol{V})\, g(\boldsymbol{B})$.

Expanding (51), we have

$$F(q, g, \boldsymbol{V}; \boldsymbol{X}, \boldsymbol{Y}) = -\tfrac{n}{2} \log |2\pi \boldsymbol{V}| - \tfrac{1}{2} \mathrm{tr}(\boldsymbol{V}^{-1} \mathrm{ERSS}) + \text{const}, \tag{52}$$

where $\mathrm{ERSS} \in \mathbb{R}^{r\times r}$ is the expected residual sum of squares, and the "const" in this expression includes additional terms in the ELBO that to not involve $\boldsymbol{V}$. Therefore, the setting of the residual covariance that maximizes the ELBO (with all other quantities fixed) is

$$\hat{\boldsymbol{V}} := \mathrm{argmax}_{\boldsymbol{V}} F(q, g, \boldsymbol{V}; \boldsymbol{X}, \boldsymbol{Y}) = \mathrm{ERSS}/n. \tag{53}$$

The expected residual sum of squares (ERSS) is

$$\mathrm{ERSS} := E_q[\boldsymbol{R}^\intercal \boldsymbol{R}], \tag{54}$$

where $\boldsymbol{R} = \boldsymbol{Y} - \boldsymbol{X}\boldsymbol{B}$ is the $n \times r$ matrix of residuals, and expectations are taken with respect to $q(\boldsymbol{B})$. After expanding and rearranging terms, the ERSS works out to

$$\mathrm{ERSS} = \bar{\boldsymbol{R}}^\intercal \bar{\boldsymbol{R}} + \sum_{j=1}^{p} \sum_{j'=1}^{p} \boldsymbol{x}_j^\intercal \boldsymbol{x}_{j'} \mathrm{Cov}_q(\boldsymbol{b}_j, \boldsymbol{b}_{j'}) \tag{55}$$

where $\bar{\boldsymbol{R}} := E_q[\boldsymbol{Y} - \boldsymbol{X}\bar{\boldsymbol{B}}] = \boldsymbol{Y} - \boldsymbol{X}\bar{\boldsymbol{B}}$ is the posterior mean residual matrix with respect to $q(\boldsymbol{B})$, $\bar{\boldsymbol{B}} := E_q[\boldsymbol{B}]$ is the posterior mean of the coefficients $\boldsymbol{B}$ with respect to $q(\boldsymbol{B})$, $\boldsymbol{x}_j = (x_{1j}, \ldots, x_{nj})^\intercal$ denotes the $j$th column of $\boldsymbol{X}$, and $\mathrm{Cov}_q(\boldsymbol{b}_j, \boldsymbol{b}_{j'}) \in \mathbb{R}^{r \times r}$ is the covariance between $\boldsymbol{b}_j$ and $\boldsymbol{b}_{j'}$, again taken with respect to $q(\boldsymbol{B})$. Notice that these expressions do not make any special assumptions about the form of $q(\boldsymbol{B})$, and so they are valid for any $q(\boldsymbol{B})$.

The solution (53) can be viewed as an M-step update in an EM algorithm for maximum-likelihood estimation of $\boldsymbol{V}$ in which expectations (computed in the E-step) are computed approximately (Blei et al., 2003, 2017).

#### G.2. Computing approximate posteriors for $\boldsymbol{B}$

In this section, we derive posterior computations for $\boldsymbol{B}$ with the assumption that the prior, $g$, is a mixture of multivariate normals (42), and with the assumption that $q$ factorizes over the variables $j$. This is summarized by the following proposition.

Proposition 5. *Suppose that the prior on the regression coefficients, $g$, is a mixture of multivariate normals* (42). *Further, we restrict $\mathcal{Q}$ to be the set of posterior distributions on $\boldsymbol{B}$ that factorize over the variables $j$, or equivalently over the columns of $\boldsymbol{B}$,*

$$q(\boldsymbol{B}) = \prod_{j=1}^{p} q_j(\boldsymbol{b}_j). \tag{56}$$

*Defined in this way, each factor $q_j(\boldsymbol{b}_j)$ is also a marginal posterior. Define a function,* BMSR-mix*, that returns the posterior distribution of $\boldsymbol{b}$ under the Bayesian multivariate regression model with a mixture-of-normals prior (see Sec. F.4). Since the posterior distribution is uniquely determined by the posterior assignment probabilities $w_{1,k}$, the posterior means $\boldsymbol{b}_{1,k}$ and the covariances $\boldsymbol{S}_{1,k}$, we write this as*

$$\text{BMSR-mix}(\boldsymbol{x}, \boldsymbol{Y}, \boldsymbol{V}, \mathscr{S}_0, \boldsymbol{w}_0) := (w_{1,1}, \ldots, w_{1,K}, \boldsymbol{b}_{1,1}, \ldots, \boldsymbol{b}_{1,K}, \boldsymbol{S}_{1,1}, \ldots, \boldsymbol{S}_{1,K}). \tag{57}$$

*Then the update for $q_j(\boldsymbol{b}_j)$—that is, the $q_j(\boldsymbol{b}_j)$ that maximizes the ELBO* (50) *while $g, \boldsymbol{V}$ and the other factors $q_{j'}(\boldsymbol{b}_{j'}), j' \neq j$ are fixed—has a closed-form solution:*

$$\begin{aligned} \hat{q}_j &:= \mathrm{argmax}_{q_j}\, F(q, g, \boldsymbol{V}; \boldsymbol{X}, \boldsymbol{Y}) \\ &= \text{BMSR-mix}(\boldsymbol{x}_j, \bar{\boldsymbol{R}}_j, \boldsymbol{V}; \mathscr{S}_0, \boldsymbol{w}_0), \end{aligned} \tag{58}$$

*in which we define $\bar{\boldsymbol{R}}_j := E_q[\boldsymbol{Y} - \sum_{j' \neq j} \boldsymbol{x}_{j'} \boldsymbol{b}_{j'}^\intercal] = \boldsymbol{Y} - \sum_{j' \neq j} \boldsymbol{x}_{j'} \bar{\boldsymbol{b}}_{j'}^\intercal$ as the $n \times r$ matrix of residuals that ignore the $j$th variable.*

Corollary 1. *A direct corollary of Proposition 5 is that the optimal $q(\boldsymbol{B})$ restricted according to* (56) *is a product of factors in which the individual factors are mixtures of multivariate normals, which we write as*

$$q_j(\boldsymbol{b}_j) = \sum_{k=1}^{K} w_{1,k}^{(j)} N_r(\boldsymbol{b}_j; \boldsymbol{b}_{1,k}^{(j)}, \boldsymbol{S}_{1,k}^{(j)}). \tag{59}$$

In summary, we have imposed a conditional independence assumption (56) on the posterior of $\boldsymbol{B}$, namely that $\boldsymbol{b}_j$ is independent of the other coefficients. Imposing this conditional independence assumption naturally leads to a coordinatewise updating procedure in which $q_j(\boldsymbol{b}_j)$ is optimized while the remaining coordinates $q_{j'}(\boldsymbol{b}_{j'})$, $j' \neq j$, are fixed, and each of the coordinatewise updates has a simple, closed-form solution (58). A corollary is that the optimal $q_j(\boldsymbol{b}_j)$ over all possible posterior distributions on $\boldsymbol{b}_j$ is a mixture of multivariate normals.

The proof of Proposition 5 is sketched in the next two subsections.

#### G.2.1. *Special case of* $p = 1$

We have already established that posterior computations are tractable for the special case of one variable when $g$ is a mixture-of-normals prior; this is Bayesian multivariate simple regression with a mixture-of-normals prior, and the posterior quantities were derived in Sec. F.4. In particular, for

$$g(\boldsymbol{b}) = \sum_{k=1}^{K} w_{0,k} N_r(\boldsymbol{0}, \boldsymbol{S}_{0,k}), \tag{60}$$

the posterior distribution of $\boldsymbol{b}$ is

$$p(\boldsymbol{b} \mid \boldsymbol{x}, \boldsymbol{Y}, \boldsymbol{V}, g) = \sum_{k=1}^{K} w_{1,k} N_r(\boldsymbol{b}_{1,k}, \boldsymbol{S}_{1,k}). \tag{61}$$

The ELBO for the *mr.mash* model with $p = 1$ variable (dropping the unneeded $j$ subscripts) is

$$\begin{aligned} F^{(p=1)}(q, g, \boldsymbol{V}; \boldsymbol{x}, \boldsymbol{Y}) &= E_q[\log p(\boldsymbol{Y} \mid \boldsymbol{x}, \boldsymbol{V}, \boldsymbol{b})] - D_{\mathrm{KL}}(q(\boldsymbol{b}) \,\|\, g(\boldsymbol{b})) \\ &= -\tfrac{n}{2}\log|2\pi\boldsymbol{V}| - \tfrac{1}{2}\mathrm{tr}(\boldsymbol{V}^{-1}\mathrm{ERSS}) - D_{\mathrm{KL}}(q(\boldsymbol{b}) \,\|\, g(\boldsymbol{b})), \end{aligned} \tag{62}$$

where

$$\begin{aligned} \mathrm{ERSS} &= E_q[(\boldsymbol{Y} - \boldsymbol{x}\boldsymbol{b}^\intercal)^\intercal(\boldsymbol{Y} - \boldsymbol{x}\boldsymbol{b}^\intercal)] \\ &= (\boldsymbol{Y} - \boldsymbol{x}\bar{\boldsymbol{b}}^\intercal)^\intercal(\boldsymbol{Y} - \boldsymbol{x}\bar{\boldsymbol{b}}^\intercal) + \boldsymbol{x}^\intercal\boldsymbol{x}\,\mathrm{Var}_q[\boldsymbol{b}], \end{aligned} \tag{63}$$

and where $\bar{\boldsymbol{b}} := E_q[\boldsymbol{b}] = \boldsymbol{b}_1^{\mathsf{mix}}$ is the posterior mean of the coefficients $\boldsymbol{b}$ with respect to the distribution $q(\boldsymbol{b})$ [see eq. 45], and $\mathrm{Var}_q[\boldsymbol{b}] = \boldsymbol{S}_1^{\mathsf{mix}}$ is the covariance of $\boldsymbol{b}$, again with respect to $q(\boldsymbol{b})$ [see eq. 46].

When $\mathcal{Q}$ is the set of all valid posterior distributions on $\mathbb{R}^r$, the variational distribution $q \in \mathcal{Q}$ maximizing the ELBO (62), $\hat{q} := \mathrm{argmax}_q\, F^{(p=1)}(q, g, \boldsymbol{V}; \boldsymbol{x}, \boldsymbol{Y})$, is the true posterior, $\hat{q}(\boldsymbol{b}) = p(\boldsymbol{b} \mid \boldsymbol{x}, \boldsymbol{Y}, \boldsymbol{V}, g)$. At $\hat{q}$, the ELBO (62) is equal to the marginal log-likelihood; that is, $F^{(p=1)}(\hat{q}, g, \boldsymbol{V}; \boldsymbol{x}, \boldsymbol{Y}) = \log p(\boldsymbol{Y} \mid \boldsymbol{x}, \boldsymbol{V}, g)$.

#### G.2.2. *Updating* $q_j(\boldsymbol{b}_j)$

To allow for posterior computations that are tractable, we restricted $\mathcal{Q}$ to the set of posterior distributions that factorize over the variables $j$; see eq. 56. With this approximation, we can now divide and conquer; we consider the problem of finding a $q_j(\boldsymbol{b}_j)$ that maximizes the ELBO (50) while the remaining factors are fixed.

Expanding only the terms involving $q_j$, the ELBO reduces to

$$\begin{aligned} F(q, g, \boldsymbol{V}; \boldsymbol{X}, \boldsymbol{Y}) &= -\tfrac{n}{2}\log|2\pi\boldsymbol{V}| - \tfrac{1}{2}\mathrm{tr}(\boldsymbol{V}^{-1}\mathrm{ERSS}) - D_{\mathrm{KL}}(q_j(\boldsymbol{b}_j) \,\|\, g(\boldsymbol{b}_j)) + \mathrm{const} \\ &= -\tfrac{n}{2}\log|2\pi\boldsymbol{V}| - \tfrac{1}{2}\mathrm{tr}[\boldsymbol{V}^{-1}(\boldsymbol{Y} - \textstyle\sum_{j=1}^{p}\boldsymbol{x}_j\bar{\boldsymbol{b}}_j^\intercal)^\intercal(\boldsymbol{Y} - \textstyle\sum_{j=1}^{p}\boldsymbol{x}_j\bar{\boldsymbol{b}}_j^\intercal)] \\ &\qquad - \tfrac{1}{2}\boldsymbol{x}_j^\intercal\boldsymbol{x}_j\mathrm{tr}(\boldsymbol{V}^{-1}\mathrm{Var}_q[\boldsymbol{b}_j]) - D_{\mathrm{KL}}(q_j(\boldsymbol{b}_j) \,\|\, g(\boldsymbol{b}_j)) + \mathrm{const} \\ &= F^{(p=1)}(q_j, g, \boldsymbol{V}; \boldsymbol{x}_j, \bar{\boldsymbol{R}}_j) + \mathrm{const}. \end{aligned} \tag{64}$$

where the "const" in each expression includes additional terms not involving $q_j$. The covariance terms $\mathrm{Cov}_q(\boldsymbol{b}_j, \boldsymbol{b}_{j'})$ for all $j \neq j'$ disappear from the ERSS because they are zero by assumption (56), and note that $\mathrm{Cov}_q(\boldsymbol{b}_j, \boldsymbol{b}_j) = \mathrm{Var}_q(\boldsymbol{b}_j)$. In summary, the ELBO can be rearranged to exactly match the expression for the single-variable ELBO (62) if we ignore terms not involving $q_j$.

#### G.3. Computing the ELBO

While computing the *mr.mash* ELBO (50) is not required, it is useful for monitoring progress of the model fitting, and for model comparison (e.g., comparing different *mr.mash* model fits). Here we explain how we compute the ELBO.

Expanding terms in (50), we have

$$F(q, g, \boldsymbol{V}; \boldsymbol{X}, \boldsymbol{Y}) = -\tfrac{n}{2}\log|2\pi\boldsymbol{V}| - \tfrac{1}{2}\mathrm{tr}(\boldsymbol{V}^{-1}\mathrm{ERSS}) - \textstyle\sum_{j=1}^{p} D_{\mathrm{KL}}(q_j(\boldsymbol{b}_j)\,\|\,g(\boldsymbol{b}_j)), \tag{65}$$

where the ERSS, starting from (55), simplifies to

$$\mathrm{ERSS} = \bar{\boldsymbol{R}}^\intercal\bar{\boldsymbol{R}} + \sum_{j=1}^{p}\boldsymbol{x}_j^\intercal\boldsymbol{x}_j\mathrm{Var}_q[\boldsymbol{b}_j]. \tag{66}$$

Recall, $\mathrm{Cov}_q(\boldsymbol{b}_j, \boldsymbol{b}_{j'}) = 0$ for all $j \neq j'$ when assuming the factorization (56). The means $\bar{b}_j$ and variances $\mathrm{Var}_q[\boldsymbol{b}_j]$ appearing in the ERSS are easily computed; see Sec. G.2.

The remaining terms in the ELBO (65) to work out are the KL-divergences. We show next that it is most convenient to compute each KL-divergence $D_{\mathrm{KL}}(q_j(\boldsymbol{b}_j)\,\|\,g(\boldsymbol{b}_j))$ when $q(\boldsymbol{b}_j)$ is updated.

##### G.3.1. Kullback-Leibler divergence for special case of $p = 1$

Starting from (62), the KL-divergence for the *mr.mash* model with one variable is

$$\begin{aligned} D_{\mathrm{KL}}(q(\boldsymbol{b})\,\|\,g(\boldsymbol{b})) &= E_q[\log p(\boldsymbol{Y} \mid \boldsymbol{x}, \boldsymbol{V}, \boldsymbol{b})] - F^{(p=1)}(q, g, \boldsymbol{V}; \boldsymbol{x}, \boldsymbol{Y}) \\ &= -\tfrac{n}{2}\log|2\pi\boldsymbol{V}| - \tfrac{1}{2}\mathrm{tr}(\boldsymbol{V}^{-1}\mathrm{ERSS}) - F^{(p=1)}(q, g, \boldsymbol{V}; \boldsymbol{x}, \boldsymbol{Y}), \end{aligned} \tag{67}$$

in which the ERSS for the single-variant *mr.mash* model is given in (63). Recall, the optimal variational distribution is equal to the true posterior, $\hat{q}(\boldsymbol{b}) = p(\boldsymbol{b} \mid \boldsymbol{x}, \boldsymbol{Y}, \boldsymbol{V}, g)$, and when the optimal variational distribution is obtained, the ELBO is equal to the marginal log-likelihood. Therefore, we have

$$\begin{aligned} D_{\mathrm{KL}}(\hat{q}(\boldsymbol{b})\,\|\,g(\boldsymbol{b})) &= -\tfrac{n}{2}\log|2\pi\boldsymbol{V}| - \tfrac{1}{2}\mathrm{tr}(\boldsymbol{V}^{-1}\mathrm{ERSS}) - \log p(\boldsymbol{Y} \mid \boldsymbol{x}, \boldsymbol{b}, \boldsymbol{V}, g) \\ &= -\tfrac{n}{2}\log|2\pi\boldsymbol{V}| - \tfrac{1}{2}\mathrm{tr}(\boldsymbol{V}^{-1}\mathrm{ERSS}) \\ &\quad - \log\mathrm{BF}^{\mathsf{mix}}(\boldsymbol{x}, \boldsymbol{Y}, \boldsymbol{V}, \mathscr{S}_0, \boldsymbol{w}_0) - \log p(\boldsymbol{Y} \mid \boldsymbol{x}, \boldsymbol{b} = \boldsymbol{0}, \boldsymbol{V}) \\ &= -\tfrac{1}{2}\mathrm{tr}[\boldsymbol{V}^{-1}(\mathrm{ERSS} - \boldsymbol{Y}^\intercal\boldsymbol{Y})] - \log\mathrm{BF}^{\mathsf{mix}}(\boldsymbol{x}, \boldsymbol{Y}, \boldsymbol{V}, \mathscr{S}_0, \boldsymbol{w}_0), \end{aligned} \tag{68}$$

where the the Bayes factor is given in (44). Finally, to arrive at the desired KL-divergences in (67), $D_{\mathrm{KL}}(\hat{q}_j(\boldsymbol{b}_j)\,\|\,g(\boldsymbol{b}_j))$, we set $\boldsymbol{x}$ to $\boldsymbol{x}_j$, and substitute $\bar{\boldsymbol{R}}_j$ for $\boldsymbol{Y}$ in (68).

#### G.4. Estimating $g$

If we assume that the prior covariances are $\boldsymbol{S}_{0,k}$ are known, or have been previously estimated using some other statistical procedure, estimating the mixture-of-normals prior $g$ reduces to estimating the mixture weights $\boldsymbol{w}_0$. Fitting the mixture weights by directly optimizing the ELBO (50) is not practical because it involves (intractable) expectations of sums inside logarithms, $E_q[\log g(\boldsymbol{b}_j)]$. Therefore, to derive a practical approach, we use an augmented *mr.mash* model that is equivalent to the original *mr.mash* model after integrating, or averaging, over the newly introduced variables. We then optimize the ELBO for this augmented model. Although this approach is optimizing a different objective, the updates for $\boldsymbol{V}$ and $q(\boldsymbol{B})$ would remain unchanged if we derived them from the augmented model, so we can interpret the above updates as also optimizing the ELBO for the augmented model.

The augmented *mr.mash* model is

$$\begin{aligned} \boldsymbol{Y} &\sim MN_{n\times r}(\boldsymbol{X}\boldsymbol{B}, \boldsymbol{I}_n, \boldsymbol{V}) \\ p(\boldsymbol{b}_j \mid \gamma_j = k) &\sim N_r(\boldsymbol{0}, \boldsymbol{S}_{0,k}) \\ p(\gamma_j = k) &= w_{0,k}, \end{aligned} \tag{69}$$

where we have introduced latent indicator variables $\boldsymbol{\gamma} = \{\gamma_1, \ldots, \gamma_p\}$, $\gamma_j \in \{1, \ldots, K\}$. Integrating over $\boldsymbol{\gamma}$ recovers the original *mr.mash* model (48).

The ELBO for the augmented model is

$$\tilde{F}(\tilde{q}, g, \boldsymbol{V}; \boldsymbol{X}, \boldsymbol{Y}) = E_{\tilde{q}}[\log p(\boldsymbol{Y} \mid \boldsymbol{X}, \boldsymbol{V}, \boldsymbol{B})] - \textstyle\sum_{j=1}^p D_{\mathrm{KL}}(\tilde{q}_j(\boldsymbol{b}_j, \gamma_j) \,\|\, p(\boldsymbol{b}_j, \gamma_j)), \tag{70}$$

in which $\tilde{q}(\boldsymbol{B}, \boldsymbol{\gamma})$ is an approximate posterior distribution on $\boldsymbol{B}, \boldsymbol{\gamma}$, and $\tilde{q}_j(\boldsymbol{b}_j, \gamma_j)$ is the (approximate) marginal posterior for $\boldsymbol{b}_j, \gamma_j$. As before, we restrict $\tilde{q}(\boldsymbol{B}, \boldsymbol{\gamma})$ to distributions that factorize over the variables $j$:

$$\tilde{q}(\boldsymbol{B}, \boldsymbol{\gamma}) = \prod_{j=1}^p \tilde{q}_j(\boldsymbol{b}_j, \gamma_j). \tag{71}$$

To derive an update for the mixture weights $\boldsymbol{w}_0$ in the mixture-of-normals prior $g$, we note that $g$ only appears in the KL-divergence terms in (70). Expanding only the parts of the ELBO that involve $\boldsymbol{w}_0$, we have

$$\begin{aligned} \tilde{F}(\tilde{q}, g, \boldsymbol{V}; \boldsymbol{X}, \boldsymbol{Y}) &= \textstyle\sum_{j=1}^p E_{\tilde{q}_j}[\log p(\gamma_j \mid \boldsymbol{w}_0)] + \text{const}, \\ &= \textstyle\sum_{j=1}^p \sum_{k=1}^K w_{1,k}^{(j)} \log w_{0,k}, \end{aligned} \tag{72}$$

where $w_{1,k}^{(j)}$ is defined in (59), and "const" is a placeholder for additional terms that do not involve $\boldsymbol{w}_0$. Therefore, the setting of the mixture weights that maximizes the ELBO (70), with all other quantities fixed, is

$$\hat{\boldsymbol{w}}_0 := \mathrm{argmax}_{\boldsymbol{w}_0} \tilde{F}(\tilde{q}, g, \boldsymbol{V}; \boldsymbol{X}, \boldsymbol{Y}) = (\hat{w}_{0,1}, \ldots, \hat{w}_{0,K}), \tag{73}$$

where $\hat{w}_{0,k} = \frac{1}{p}\sum_{j=1}^p w_{1,k}^{(j)}$. Similar to the update for $\boldsymbol{V}$, we can view (73) as an exact M-step with an approximate E-step.

### H. Derivation of *mr.mash* variational algorithms with missing data

Here we treat the case in which one or more entries of $\boldsymbol{Y}$ are missing. Our approach involves treating the missing entries, $\boldsymbol{Y}_{\text{miss}}$, as random variables, and modeling them using the *mr.mash* model, then formulating a variational approximation with the assumption that $\boldsymbol{Y}_{\text{miss}}$ is independent of $\boldsymbol{B}$. One difference compared to the fully observed case is that we explicitly include an intercept in the multivariate regression, and treat it as a free parameter to be estimated (by maximizing the ELBO).

The *mr.mash* model with an intercept is

$$\begin{aligned} \boldsymbol{Y} &\sim MN_{n \times r}(\mathbf{1}_n \boldsymbol{b}_0^\intercal + \boldsymbol{X}\boldsymbol{B}, \boldsymbol{I}_n, \boldsymbol{V}) \\ \boldsymbol{b}_j &\overset{i.i.d.}{\sim} g, \text{ for } j = 1, \ldots, p, \end{aligned} \tag{74}$$

in which $\boldsymbol{b}_0 \in \mathbb{R}^r$ is the intercept.

Next, taking a variational inference approach, we compute an approximate posterior $q(\boldsymbol{Y}_{\text{miss}}, \boldsymbol{B})$, making the following conditional independence assumption:

$$q(\boldsymbol{Y}_{\text{miss}}, \boldsymbol{B}) = q(\boldsymbol{Y}_{\text{miss}})\, q(\boldsymbol{B}), \tag{75}$$

and the approximate posterior for $\boldsymbol{B}$ factorizes as before (see eq. 9).

Importantly, as we will see, with this conditional independence assumption we can reuse the computations from the fully-observed case. The ELBO with missing data is

$$\begin{aligned} F_{\text{miss}}(q, g, \boldsymbol{V}, \boldsymbol{b}_0; \boldsymbol{X}, \boldsymbol{Y}_{\text{obs}}) = \mathrm{E}_q[\log p(\boldsymbol{Y} \mid \boldsymbol{X}, \boldsymbol{V}, \boldsymbol{b}_0, \boldsymbol{B})] - \mathrm{E}_q[\log q(\boldsymbol{Y}_{\text{miss}})] \\ - \sum_{j=1}^p D_{\mathrm{KL}}(q_j(\boldsymbol{b}_j) \,\|\, g(\boldsymbol{b}_j)), \end{aligned} \tag{76}$$

where $\boldsymbol{Y}_{\text{obs}}$ denotes the observed responses.

In the expressions below, we use $\bar{\boldsymbol{Y}}$ to denote the "imputed" $\boldsymbol{Y}$ in which missing entries are replaced by their approximate posterior means $E_q[\boldsymbol{Y}_{\text{miss}}]$, and observed entries are copied over from $\boldsymbol{Y}_{\text{obs}}$. And $\boldsymbol{Y}$ denotes the combination of the observed and missing values, in which the missing values are treated as random variables.

#### H.1. Estimating $\boldsymbol{b}_0$

First, we focus on the parts of the ELBO (76) that involve $\boldsymbol{b}_0$. We have,

$$F_{\text{miss}}(q, g, \boldsymbol{V}, \boldsymbol{b}_0; \boldsymbol{X}, \boldsymbol{Y}_{\text{obs}}) = -\tfrac{n}{2}\boldsymbol{b}_0^\intercal \boldsymbol{V}^{-1}\boldsymbol{b}_0 - \tfrac{1}{n}(\bar{\boldsymbol{m}}_y - \bar{\boldsymbol{B}}^\intercal \boldsymbol{m}_x)\boldsymbol{V}^{-1}\boldsymbol{b}_0 + \text{const}, \tag{77}$$

in which $\boldsymbol{m}_x = \boldsymbol{X}^\intercal \mathbf{1}_n/n$ is the vector of length $p$ containing the column means of $\boldsymbol{X}$, $\bar{\boldsymbol{m}}_y = \bar{\boldsymbol{Y}}^\intercal \mathbf{1}/n$ is the vector of length $r$ containing the column means of $\bar{\boldsymbol{Y}}$, and the "const" includes the additional terms in the ELBO that do not involve $\boldsymbol{b}_0$. The maximizer is therefore at

$$\hat{\boldsymbol{b}}_0 = \bar{\boldsymbol{m}}_y - \bar{\boldsymbol{B}}^\intercal \boldsymbol{m}_x. \tag{78}$$

#### H.2. Computing approximate posteriors for $\boldsymbol{B}$

Using (78), one can show that the following identity holds:

$$\max_{\boldsymbol{b}_0} F_{\text{miss}}(q, g, \boldsymbol{V}, \boldsymbol{b}_0; \boldsymbol{X}, \boldsymbol{Y}_{\text{obs}}) = F(q, g, \boldsymbol{V}; \tilde{\boldsymbol{X}}, \tilde{\boldsymbol{Y}}) + \text{const}, \tag{79}$$

in which $\tilde{\boldsymbol{X}} := \boldsymbol{X} - \mathbf{1}_n \boldsymbol{m}_x^\intercal$ is the column-centered data matrix, $\tilde{\boldsymbol{Y}} := \bar{\boldsymbol{Y}} - \mathbf{1}_n \bar{\boldsymbol{m}}_y^\intercal$ is the column-centered imputed response matrix, and the "const" includes terms that do not involve $\boldsymbol{B}$, $\boldsymbol{V}$ or the prior, $g$. Therefore, the problem of optimizing $q(\boldsymbol{B})$ (as well as estimating $\boldsymbol{V}$, $\boldsymbol{w}_0$), with $q(\boldsymbol{Y}_{\text{miss}})$ kept fixed, reduces to the problem of fitting a *mr.mash* model in the fully-observed setting, in which $\boldsymbol{X}$ is replaced with the column-centered matrix $\tilde{\boldsymbol{X}}$, and $\boldsymbol{Y}$ is replaced with the "imputed" matrix $\tilde{\boldsymbol{Y}}$ that has also been column-centered. So we can reuse the computations for the fully-observed case to implement model fitting for the missing-data case.

#### H.3. Computing approximate posteriors for $\boldsymbol{Y}_{\text{miss}}$

Let $\text{miss}(i) \subset \{1, \ldots, r\}$ be the subset of dimensions that have missing values in the $i$th row of $\boldsymbol{Y}$, and let the remaining (observed) dimensions be denoted by $\text{obs}(i)$. Fixing $q(\boldsymbol{B})$ and the parameters of the *mr.mash* model, it can be shown that the approximate posterior for the $i$th row of $\boldsymbol{Y}_{\text{miss}}$ that maximizes the ELBO,

$$\hat{q}(\boldsymbol{y}_{i,j}) := \text{argmax}_{q(\boldsymbol{y}_{i,j})} F_{\text{miss}}(q, g, \boldsymbol{V}, \boldsymbol{b}_0; \boldsymbol{X}, \boldsymbol{Y}_{\text{obs}}), \tag{80}$$

in which $j = \text{miss}(i)$, is multivariate normal with mean and covariance

$$\text{E}[\boldsymbol{y}_{i,j}] = \hat{\boldsymbol{y}}_{i,j} - \boldsymbol{\Lambda}_{j,j}^{-1}\boldsymbol{\Lambda}_{j,j'}(\boldsymbol{y}_{i,j'} - \hat{\boldsymbol{y}}_{i,j'}) \tag{81}$$

$$\text{Var}[\boldsymbol{y}_{i,j}] = \boldsymbol{\Lambda}_{jj}^{-1}, \tag{82}$$

such that $j' = \text{obs}(i)$, $\boldsymbol{\Lambda} := \boldsymbol{V}^{-1}$, and $\hat{\boldsymbol{y}}_i$ denotes the fitted values in the $i$th row,

$$\hat{\boldsymbol{y}}_i = \boldsymbol{b}_0 + \bar{\boldsymbol{B}}^\intercal \boldsymbol{x}_i. \tag{83}$$

#### H.4. Computing the ELBO

We can also reuse the expressions for the ELBO in the fully-observed case to compute the ELBO for the missing-data case. Assuming $\boldsymbol{b}_0$ satisfies (78), the missing-data ELBO (76) can be rewritten as

$$F_{\text{miss}}(q, g, \boldsymbol{V}, \boldsymbol{b}_0; \boldsymbol{X}, \boldsymbol{Y}_{\text{obs}}) = F(q, g; \boldsymbol{V}, \tilde{\boldsymbol{X}}, \tilde{\boldsymbol{Y}}) - \frac{1}{2}\sum_{i=1}^{n} \text{E}_q[(\boldsymbol{y}_i - \bar{\boldsymbol{y}}_i)^\intercal \boldsymbol{V}^{-1}(\boldsymbol{y}_i - \bar{\boldsymbol{y}}_i)] - \text{E}_q[\log q(\boldsymbol{Y}_{\text{miss}})], \tag{84}$$

in which $\boldsymbol{y}_i$ is the $i$th row of $\boldsymbol{Y}$ and $\bar{\boldsymbol{y}}_i$ is the $i$th row of $\bar{\boldsymbol{Y}}$.

### I. Implementation notes

Although BMSR-mix outputs all the statistics needed to fully reconstruct the posterior distribution, in practice we do not need to keep track of all these statistics. In particular, if both $g$ and $\boldsymbol{V}$ are not estimated, we only need to keep track of the marginal posterior means (45). For updating $\boldsymbol{V}$, we also need to keep track of the marginal posterior variances (46). If the mixture weights in the prior $g$ are also updated, we also need to keep track of the posterior assignment probabilities $w_{1,k}^{(j)}$ (or sums of these probabilities).

The posterior computations described above are repeated many, many times, so performing these computations efficiently is key to an efficient algorithm for fitting *mr.mash* models. Our general approach is to avoid unnecessarily computation of all inverses, (Cholesky) factorizations or determinants of $r \times r$ matrices that are used to compute the key posterior quantities and the ELBOs by precomputing these quantities when possible. (One has to be careful to update these precomputed quantities whenever the relevant model parameters are also updated.)

The special case in which $\boldsymbol{X}$ is standardized so that all the columns have unit variance, there are many computations that can be reused. For example, in this special case the posterior covariance from the Bayesian multivariate simple regression with a normal prior (Sec. F.4) is the same for all the variables $j = 1, 2, \ldots, p$, and therefore posterior covariances for all the mixture components can be precomputed and reused.

Even when $\boldsymbol{X}$ is not standardized, posterior computations can still be sped up by precomputing Cholesky factorizations and eigendecompositions of quantities involving $\boldsymbol{V}$ and $\boldsymbol{S}_{0,k}$.

### J. Proof of Proposition 3

We first show the proof from the maximum-likelihood estimation perspective. We treat $\boldsymbol{\mu}$ as a free parameter to be optimized. The log-likelihood for $\boldsymbol{\mu}$ and $\boldsymbol{b}$ is

$$\log \ell(\boldsymbol{\mu}, \boldsymbol{b}; \boldsymbol{x}, \boldsymbol{Y}, \boldsymbol{V}) = -\tfrac{1}{2}\mathrm{tr}[\boldsymbol{V}^{-1}(\boldsymbol{Y} - \boldsymbol{1}\boldsymbol{\mu}^\intercal - \boldsymbol{x}\boldsymbol{b}^\intercal)^\intercal(\boldsymbol{Y} - \boldsymbol{1}\boldsymbol{\mu}^\intercal - \boldsymbol{x}\boldsymbol{b}^\intercal)] + \mathrm{const},$$

in which the “const” includes additional terms that do not involve $\boldsymbol{\mu}$ or $\boldsymbol{b}$. Differentiating with respect to $\boldsymbol{\mu}$, setting the partial derivatives to zero, and solving for $\boldsymbol{\mu}$ yields

$$\boldsymbol{V}^{-1}\boldsymbol{\mu} = \boldsymbol{V}^{-1}(\bar{\boldsymbol{y}} - \bar{x}\boldsymbol{b}),$$

and so

$$\hat{\boldsymbol{\mu}} = \bar{\boldsymbol{y}} - \bar{x}\boldsymbol{b}.$$

The profile likelihood for $\boldsymbol{b}$ is therefore

$$\begin{aligned}
\ell^*(\boldsymbol{b}; \boldsymbol{x}, \boldsymbol{Y}, \boldsymbol{V}) &:= \max_{\boldsymbol{\mu}} \ell(\boldsymbol{\mu}, \boldsymbol{b}; \boldsymbol{x}, \boldsymbol{Y}, \boldsymbol{V}) \\
&= |2\pi\boldsymbol{V}|^{-n/2} \exp\{-\tfrac{1}{2}\mathrm{tr}[\boldsymbol{V}^{-1}(\boldsymbol{Y} - \boldsymbol{1}\hat{\boldsymbol{\mu}}^\intercal - \boldsymbol{x}\boldsymbol{b}^\intercal)^\intercal(\boldsymbol{Y} - \boldsymbol{1}\hat{\boldsymbol{\mu}}^\intercal - \boldsymbol{x}\boldsymbol{b}^\intercal)]\} \\
&= |2\pi\boldsymbol{V}|^{-n/2} \exp\{-\tfrac{1}{2}\mathrm{tr}[\boldsymbol{V}^{-1}((\boldsymbol{Y} - \boldsymbol{1}\bar{\boldsymbol{y}}^\intercal) - (\boldsymbol{x} - \bar{x}\boldsymbol{1})\boldsymbol{b}^\intercal)^\intercal((\boldsymbol{Y} - \boldsymbol{1}\bar{\boldsymbol{y}}^\intercal) - (\boldsymbol{x} - \bar{x}\boldsymbol{1})\boldsymbol{b}^\intercal)]\} \\
&= \ell(\boldsymbol{b}; \tilde{\boldsymbol{x}}, \tilde{\boldsymbol{Y}}, \boldsymbol{V}).
\end{aligned}$$

The conditional posterior for $\boldsymbol{\mu}$ given $\boldsymbol{b}$ is

$$\begin{aligned}
p(\boldsymbol{\mu} \mid \boldsymbol{x}, \boldsymbol{Y}, \boldsymbol{V}, \boldsymbol{S}_{0\mu}, \boldsymbol{b}) \propto \exp\{-\tfrac{1}{2}(\boldsymbol{\mu} - n(\boldsymbol{S}_{0\mu}^{-1} + n\boldsymbol{V}^{-1})^{-1}\boldsymbol{V}^{-1}(\bar{\boldsymbol{y}} - \bar{x}\boldsymbol{b}))^\intercal(\boldsymbol{S}_{0\mu}^{-1} + n\boldsymbol{V}^{-1}) \\
\times (\boldsymbol{\mu} - n(\boldsymbol{S}_{0\mu}^{-1} + n\boldsymbol{V}^{-1})^{-1}\boldsymbol{V}^{-1}(\bar{\boldsymbol{y}} - \bar{x}\boldsymbol{b}))\},
\end{aligned}$$

which is the multivariate normal density with mean

$$n(\boldsymbol{S}_{0\mu}^{-1} + n\boldsymbol{V}^{-1})^{-1}\boldsymbol{V}^{-1}(\bar{\boldsymbol{y}} - \bar{x}\boldsymbol{b}) = \boldsymbol{S}_{1\mu}\hat{\boldsymbol{S}}_{\mu}^{-1}\hat{\boldsymbol{\mu}} = \boldsymbol{\mu}_1,$$

and covariance

$$(\boldsymbol{S}_{0\mu}^{-1} + n\boldsymbol{V}^{-1})^{-1} = (\boldsymbol{S}_{0\mu}^{-1} + \hat{\boldsymbol{S}}_{\mu}^{-1})^{-1} = \boldsymbol{S}_{1\mu}.$$

The marginal likelihood for $\boldsymbol{b}$ is

$$\begin{aligned}\ell^\star(\boldsymbol{b}; \boldsymbol{x}, \boldsymbol{Y}, \boldsymbol{V}, \boldsymbol{S}_{0\mu}) &= \int \ell(\boldsymbol{\mu}, \boldsymbol{b}; \boldsymbol{x}, \boldsymbol{Y}, \boldsymbol{V})\, p(\boldsymbol{\mu} \mid \boldsymbol{S}_{0\mu})\, d\boldsymbol{\mu} \\ &= \int |2\pi \boldsymbol{S}_{0\mu}|^{-1/2} |2\pi \boldsymbol{V}|^{-n/2} \\ &\qquad \times \exp\{-\tfrac{1}{2}\mathrm{tr}[\boldsymbol{V}^{-1}(\boldsymbol{Y} - \boldsymbol{x}\boldsymbol{b}^\intercal - \boldsymbol{1}\boldsymbol{\mu}^\intercal)^\intercal(\boldsymbol{Y} - \boldsymbol{x}\boldsymbol{b}^\intercal - \boldsymbol{1}\boldsymbol{\mu}^\intercal) + \boldsymbol{S}_{0\mu}^{-1}\boldsymbol{\mu}\boldsymbol{\mu}^\intercal]\}\, d\boldsymbol{b}_0.\end{aligned}$$

Expanding terms, we get

$$\begin{aligned}&\ell^\star(\boldsymbol{b}; \boldsymbol{x}, \boldsymbol{Y}, \boldsymbol{V}, \boldsymbol{S}_{0\mu}) \\ &\quad = |2\pi \boldsymbol{S}_{0\mu}|^{-1/2}|2\pi \boldsymbol{V}|^{-n/2} \int \exp\{-\tfrac{1}{2}\mathrm{tr}[(n\boldsymbol{V}^{-1} + \boldsymbol{S}_{0\mu}^{-1})\boldsymbol{\mu}\boldsymbol{\mu}^\intercal - 2\boldsymbol{V}^{-1}(\boldsymbol{Y} - \boldsymbol{x}\boldsymbol{b}^\intercal)^\intercal \boldsymbol{1}\boldsymbol{\mu}^\intercal \\ &\qquad\qquad + \boldsymbol{V}^{-1}(\boldsymbol{Y} - \boldsymbol{x}\boldsymbol{b}^\intercal)^\intercal(\boldsymbol{Y} - \boldsymbol{x}\boldsymbol{b}^\intercal)]\}\, d\boldsymbol{\mu} \\ &\quad = |2\pi \boldsymbol{S}_{0\mu}|^{-1/2}|2\pi \boldsymbol{V}|^{-n/2} \int \exp\{-\tfrac{1}{2}\mathrm{tr}[(n\boldsymbol{V}^{-1} + \boldsymbol{S}_{0\mu}^{-1})(\boldsymbol{\mu} - n(n\boldsymbol{V}^{-1} + \boldsymbol{S}_{0\mu}^{-1})^{-1}\boldsymbol{V}^{-1}(\bar{\boldsymbol{y}} - \bar{x}\boldsymbol{b})) \\ &\qquad\qquad \times (\boldsymbol{\mu} - n(n\boldsymbol{V}^{-1} + \boldsymbol{S}_{0\mu}^{-1})^{-1}\boldsymbol{V}^{-1}(\bar{\boldsymbol{y}} - \bar{x}\boldsymbol{b}))^\intercal \\ &\qquad\qquad - n\boldsymbol{V}^{-1}(\bar{\boldsymbol{y}} - \bar{x}\boldsymbol{b})(\bar{\boldsymbol{y}} - \bar{x}\boldsymbol{b})^\intercal n\boldsymbol{V}^{-1}(n\boldsymbol{V}^{-1} + \boldsymbol{S}_{0\mu}^{-1})^{-1} \\ &\qquad\qquad + \boldsymbol{V}^{-1}(\boldsymbol{Y} - \boldsymbol{x}\boldsymbol{b}^\intercal)^\intercal(\boldsymbol{Y} - \boldsymbol{x}\boldsymbol{b}^\intercal)]\}\, d\boldsymbol{\mu} \\ &\quad = |\boldsymbol{S}_{0\mu}|^{-1/2}|2\pi \boldsymbol{V}|^{-n/2}|n\boldsymbol{V}^{-1} + \boldsymbol{S}_{0\mu}^{-1}|^{-1/2} \exp\{-\tfrac{1}{2}\mathrm{tr}[\boldsymbol{V}^{-1}(\boldsymbol{Y} - \boldsymbol{x}\boldsymbol{b}^\intercal)^\intercal(\boldsymbol{Y} - \boldsymbol{x}\boldsymbol{b}^\intercal) \\ &\qquad\qquad - n\boldsymbol{V}^{-1}(\bar{\boldsymbol{y}} - \bar{x}\boldsymbol{b})(\bar{\boldsymbol{y}} - \bar{x}\boldsymbol{b})^\intercal n\boldsymbol{V}^{-1}(n\boldsymbol{V}^{-1} + \boldsymbol{S}_{0\mu}^{-1})^{-1}]\} \\ &\quad = |2\pi \boldsymbol{V}|^{-n/2}|\boldsymbol{S}_{0\mu}^{-1}\boldsymbol{S}_{1\mu}|^{1/2} \exp\{\tfrac{1}{2}\boldsymbol{\mu}_1^\intercal \boldsymbol{S}_{1\mu}^{-1}\boldsymbol{\mu}_1 - \tfrac{1}{2}\mathrm{tr}[\boldsymbol{V}^{-1}(\boldsymbol{Y} - \boldsymbol{x}\boldsymbol{b}^\intercal)^\intercal(\boldsymbol{Y} - \boldsymbol{x}\boldsymbol{b}^\intercal)]\}.\end{aligned}$$

When $\boldsymbol{S}_{0\mu}^{-1} \rightarrow \boldsymbol{0}$, the marginal likelihood simplifies to

$$\begin{aligned}&\ell^\star(\boldsymbol{b}; \boldsymbol{x}, \boldsymbol{Y}, \boldsymbol{V}, \boldsymbol{S}_{0\mu}) \\ &\quad = |2\pi \boldsymbol{V}|^{-n/2}|\boldsymbol{S}_{0\mu}^{-1}\hat{\boldsymbol{S}}_\mu|^{1/2} \exp\{-\tfrac{1}{2}\mathrm{tr}\left[\boldsymbol{V}^{-1}(\boldsymbol{Y} - \boldsymbol{x}\boldsymbol{b}^\intercal)^\intercal(\boldsymbol{Y} - \boldsymbol{x}\boldsymbol{b}^\intercal) - n\boldsymbol{V}^{-1}(\bar{\boldsymbol{y}} - \bar{x}\boldsymbol{b})(\bar{\boldsymbol{y}} - \bar{x}\boldsymbol{b})^\intercal\right]\} \\ &\quad = |2\pi \boldsymbol{V}|^{-n/2}|\boldsymbol{S}_{0\mu}^{-1}\hat{\boldsymbol{S}}_\mu|^{1/2} \exp\{-\tfrac{1}{2}\mathrm{tr}\left[\boldsymbol{V}^{-1}(\boldsymbol{Y} - \boldsymbol{1}\bar{\boldsymbol{y}}^\intercal - (\boldsymbol{x} - \bar{x}\boldsymbol{1})\boldsymbol{b}^\intercal)^\intercal(\boldsymbol{Y} - \boldsymbol{1}\bar{\boldsymbol{y}}^\intercal - (\boldsymbol{x} - \bar{x}\boldsymbol{1})\boldsymbol{b}^\intercal)\right]\} \\ &\quad = |\boldsymbol{S}_{0\mu}^{-1}\hat{\boldsymbol{S}}_\mu|^{1/2} \times \ell(\boldsymbol{b}; \tilde{\boldsymbol{x}}, \tilde{\boldsymbol{Y}}, \boldsymbol{V}).\end{aligned}$$

The marginal likelihood for $\boldsymbol{b}$ in the model with an intercept is therefore proportional to the likelihood for the model without an intercept (eq. 19) when $\boldsymbol{Y}$ and $\boldsymbol{x}$ are column-centered.
