## Supplementary figures and images for "A flexible empirical Bayes approach to multivariate multiple regression, and its improved accuracy in predicting multi-tissue gene expression from genotypes"

### S1 Fig

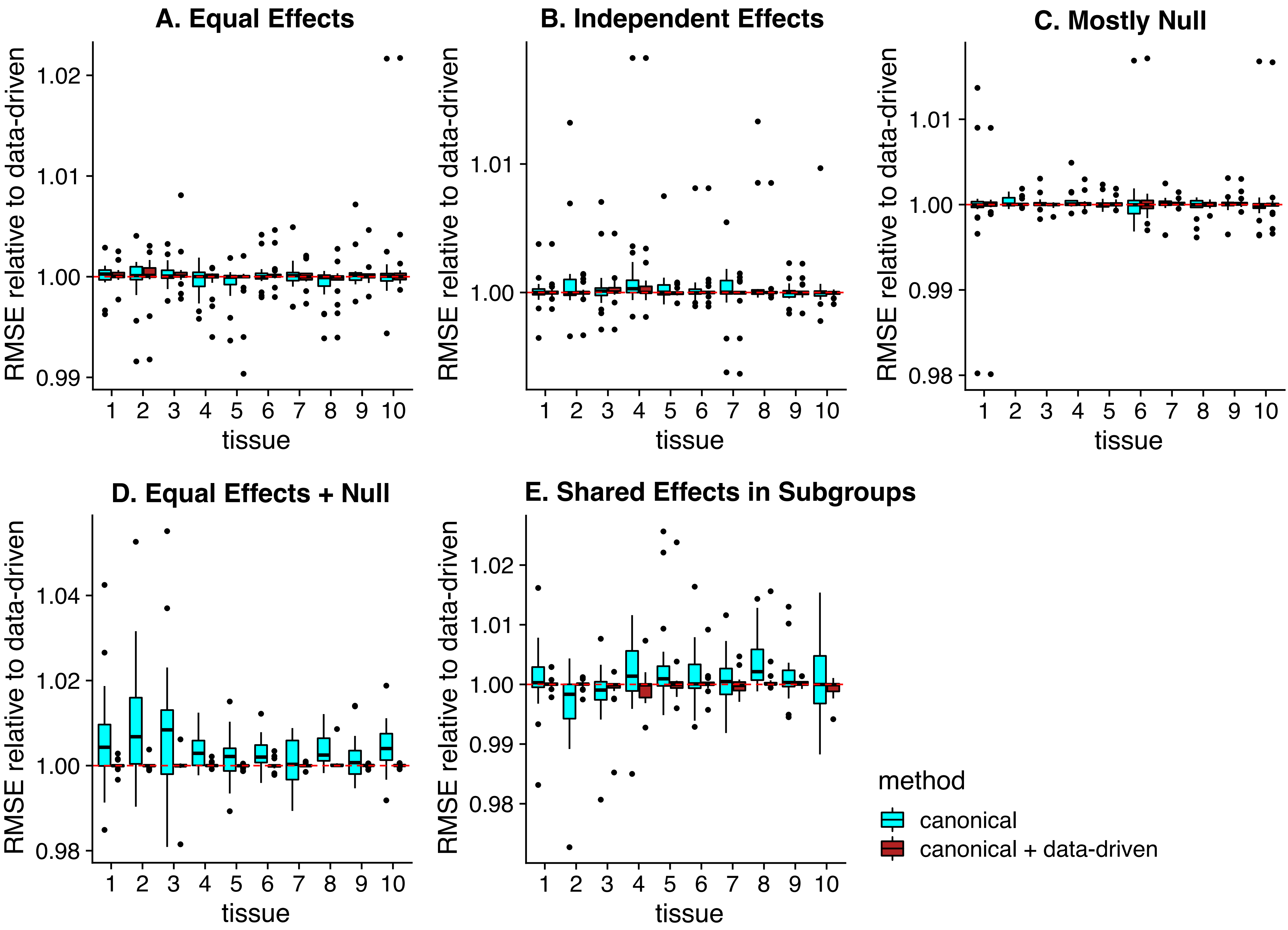

### S2 Fig

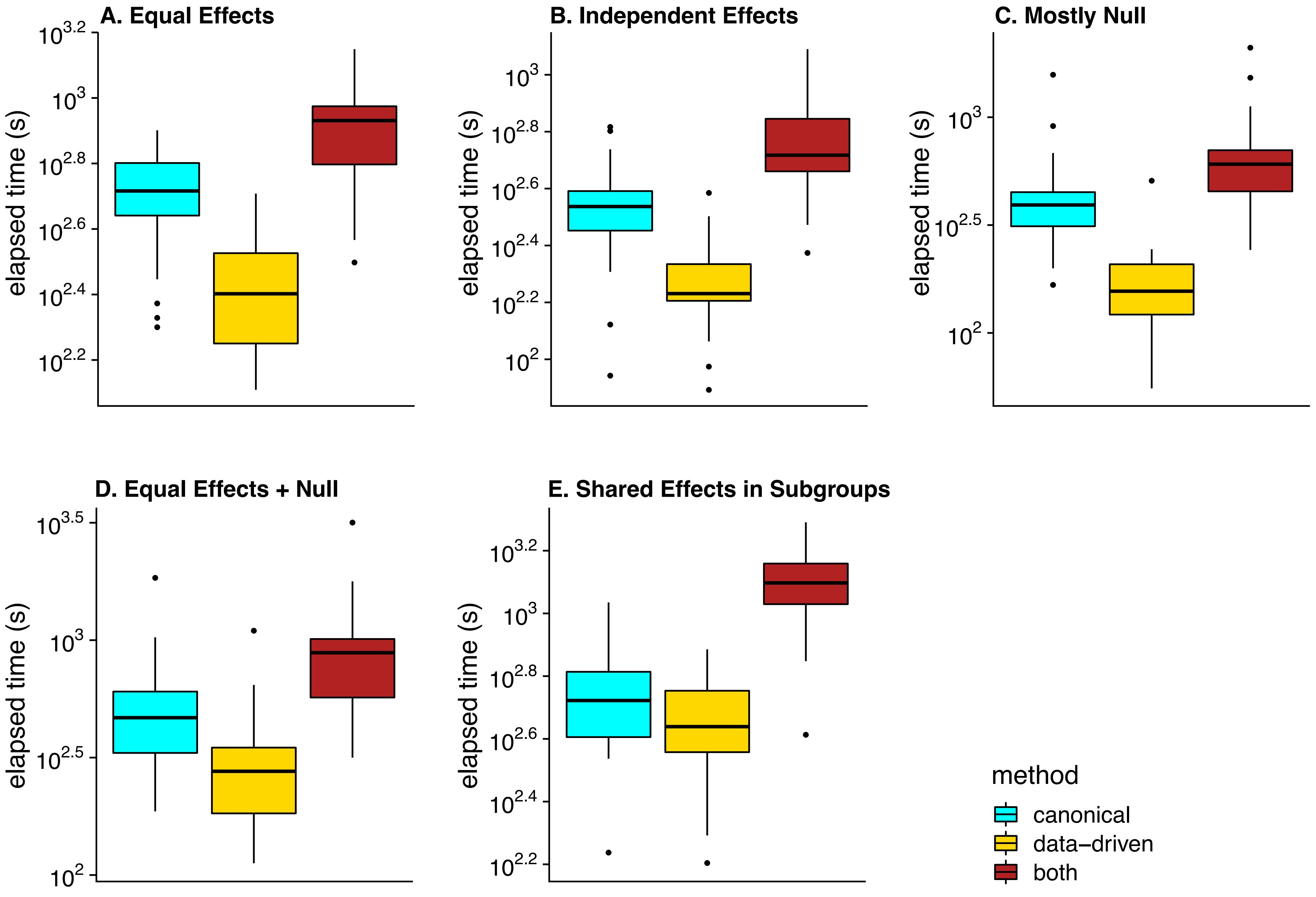

### S3 Fig

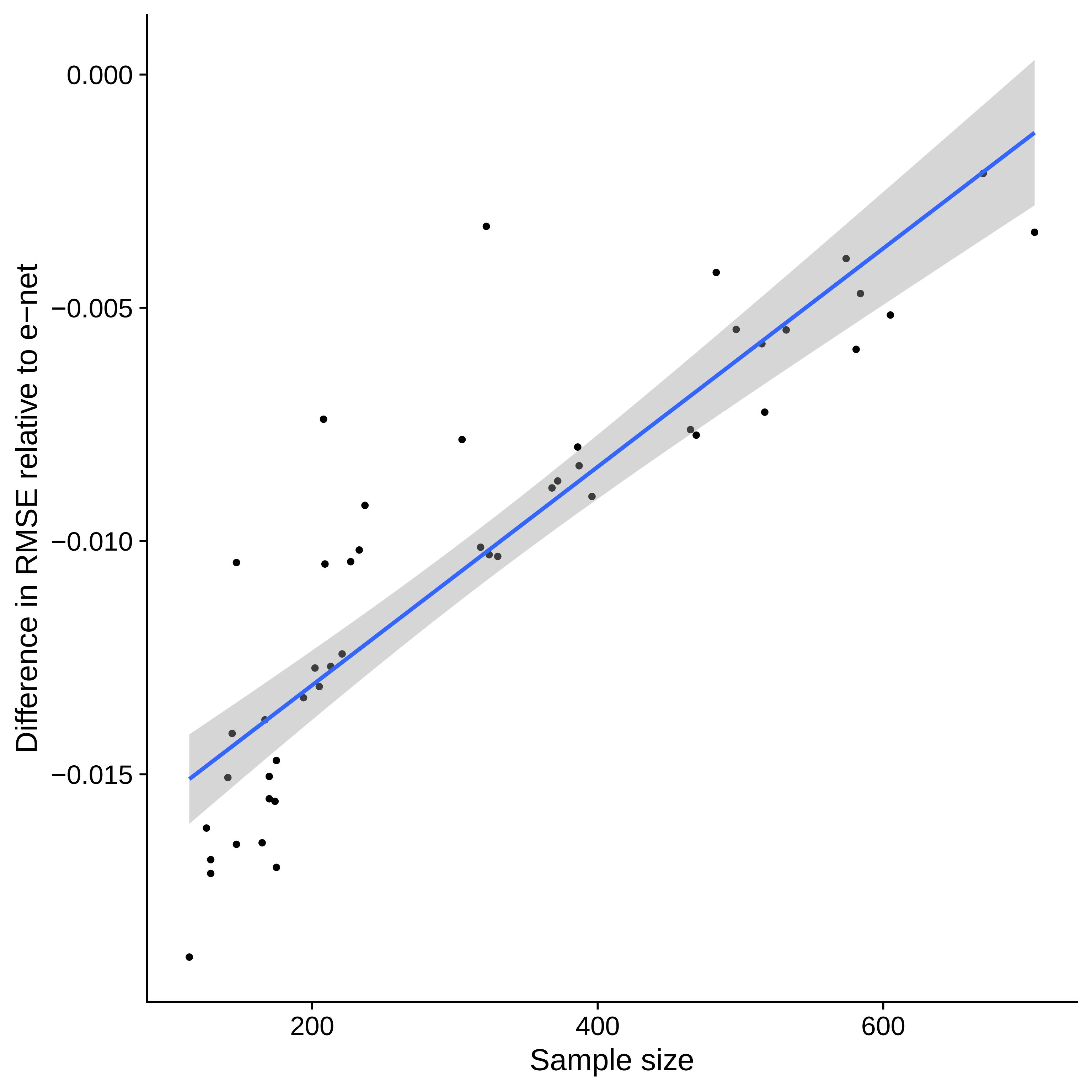

### S4 Fig

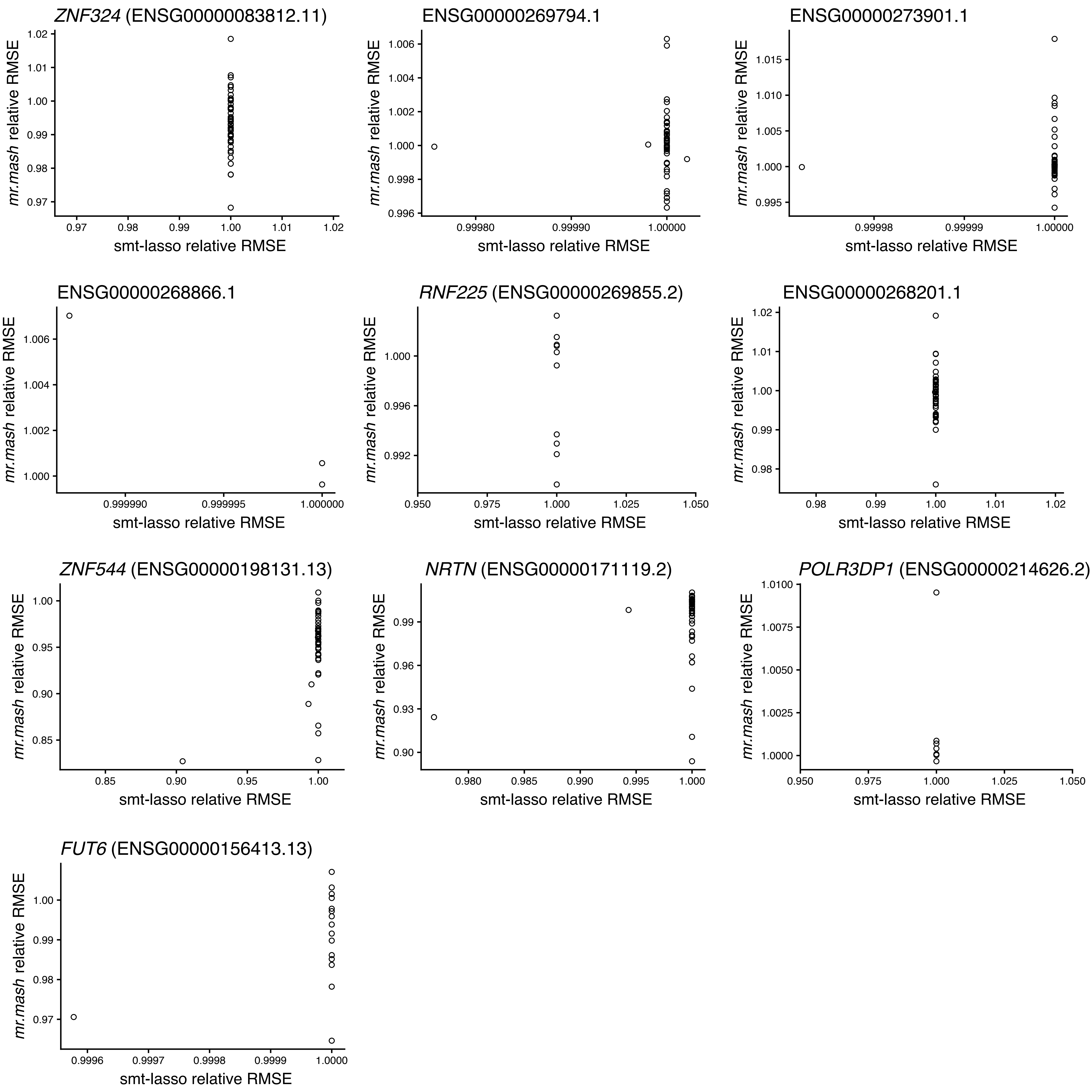

### S5 Fig

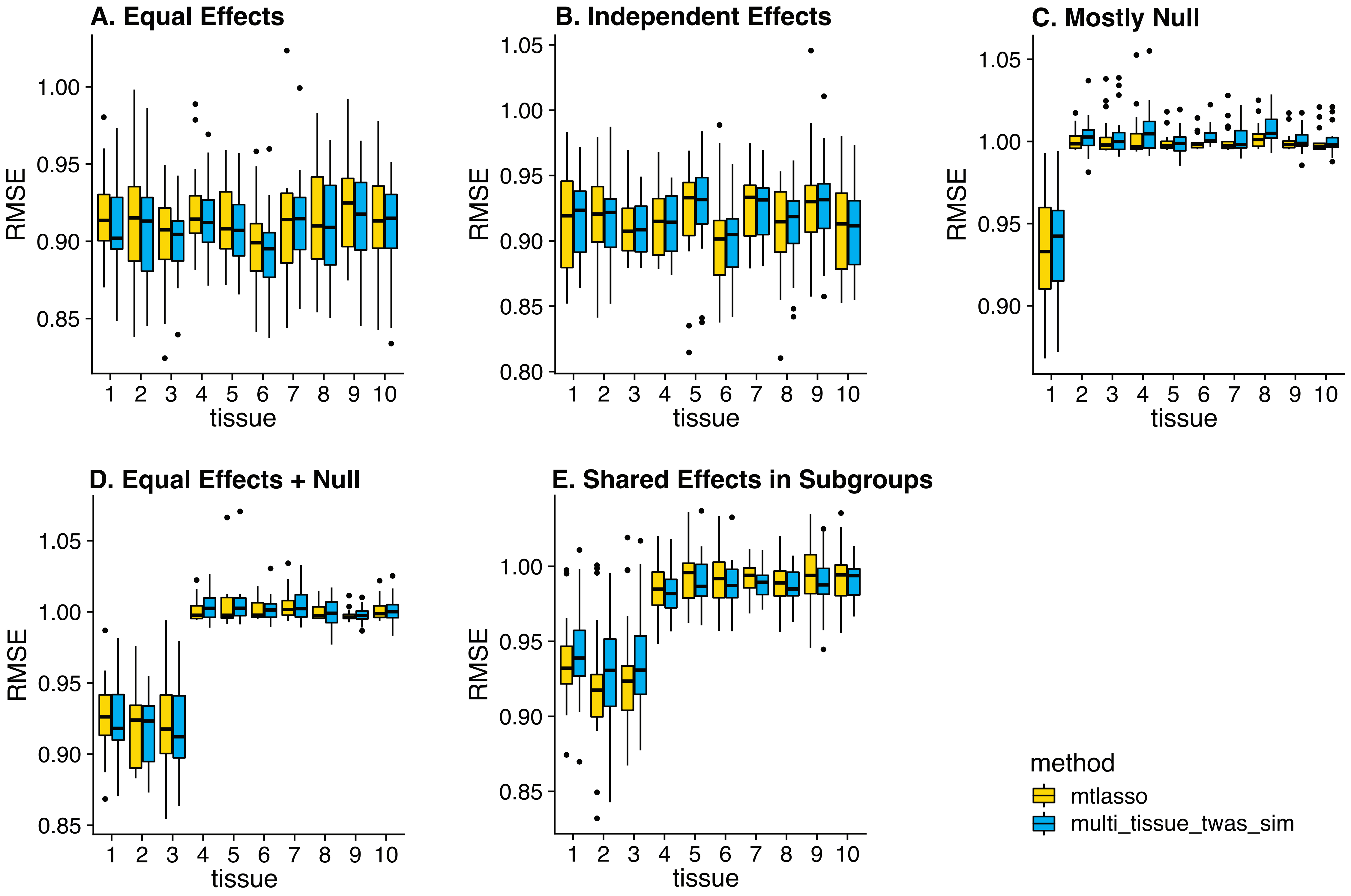

### S6 Fig

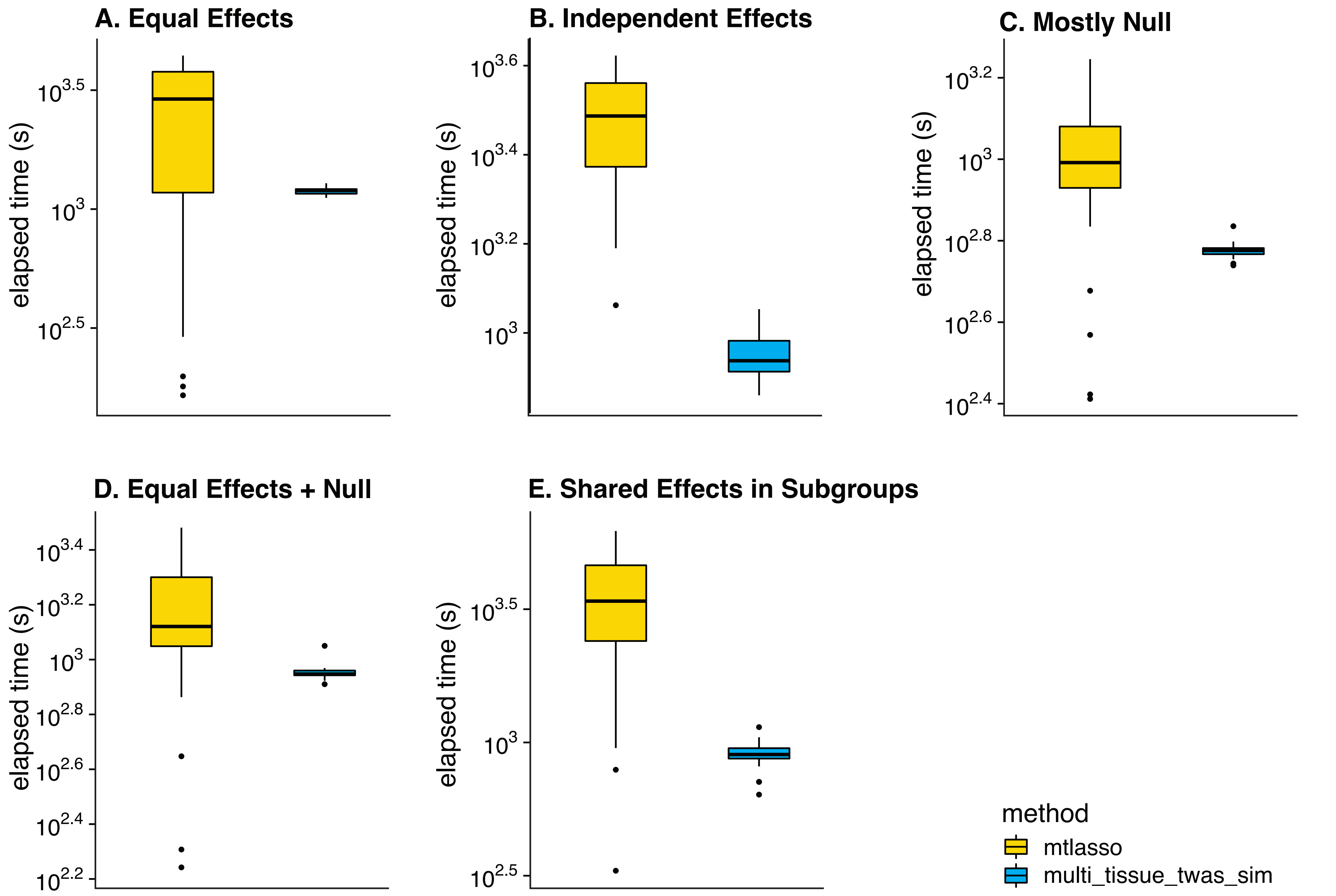
